# Postnatal maturation of putamen microstructure accompanies topographic white matter connectivity and altered circuits in autism

**DOI:** 10.64898/2026.09.09.750447

**Authors:** Vaidehi S. Natu, Christina Tyagi, Xiaoqian Yan, Sarah Tung, Emily Kubota, Seda Karakose-Akbiyik, Hua Wu, Aviv Mezer, Elior Drori, Nan Wang, Congyu Liao, Xiaozhi Cao, Kawin Setsompop, Kalanit Grill-Spector

**Author notes:** **Corresponding author:** Vaidehi S. Natu.

## Abstract

The putamen is a major hub of the basal ganglia that emerges early in gestation. However, whether its mature organization is established before birth or emerges postnatally remains unknown. Using cross-sectional and longitudinal quantitative MRI (R_1_ and R_2_*, related to tissue density and iron, respectively), and diffusion MRI, we characterized the development of putamen’s microstructure and its white matter connectivity with cortex from birth to 12 months and compared their trajectories with those in adults. Despite its prenatal emergence, the putamen undergoes substantial postnatal development. R_1_ increases from birth to 12 months, producing a prominent anterior–posterior gradient, whereas R_2_* increases primarily between age one and adulthood, producing a medial– lateral gradient. Cortico-putamen white matter connectivity is diffuse in infants but becomes topographic in adults, with anterior putamen linked to frontal cortex and posterior putamen to sensorimotor cortex. In autism spectrum disorder, this organization is largely preserved and accompanied by increased anterior putamen–prefrontal connectivity. Our findings reveal distinct spatial developmental trajectories of putamen microstructure and cortical connectivity providing a developmental framework for understanding the organization of the putamen in infancy, which has implications for assessing neurodevelopmental disorders of the basal ganglia.

**Teaser:** From birth to one year, the putamen develops distinct microstructural gradients and increasingly topographic cortical connections.

## Introduction

The putamen, a part of the dorsal striatum, is a major gateway and relay network of the basal ganglia. In adults, it is structurally organized and well-connected to prefrontal, motor, and somatosensory regions translating intentions, decisions, and rewards into coordinated motor functions and goal directed cognitive functions (*1–8*). Along the putamen’s anterior-posterior (AP, front to back) axis, functionally distinct white matter (WM) projections connect it with sensorimotor and associative cortices (*5*, *9–15*). Additionally, animal and *ex vivo* studies in adults have demonstrated distinct spatial gradients in the dorsal striatum, marked by increasing cellular density (*16*) and neurotransmitters like dopamine (*17*) along the AP axis and cellular gradients along the medial-lateral (ML) axis (*16*). A recent *in vivo* study in younger and older adults using quantitative MRI (qMRI) metrics (longitudinal relaxation rate R_1_, sensitive to tissue density and transverse relaxation rate R_2_*, sensitive to iron) (*18–21*), reveal spatial gradients with increasing R_1_ and R_2_* from the anterior to posterior putamen (*22*). However, to-date microstructural gradients and white matter connections of the putamen have only been examined in human adults and animal brains (*4*, *17*), thus, its development remains unknown.

Developmentally, the putamen emerges *in utero* in the first trimester (*23*, *24*), its neural count (*25*) and volume increase in the second trimester (*26*) as well as postnatally (*27–33*). Studies of the broader basal ganglia have reported postnatal increases in fractional anisotropy and quantitative MRI metric R_2_* (*34–36*) but *in vivo*developmental studies of the putamen have largely focused on volumetric growth. These studies revealed that atypical putamen volume is associated with neurodevelopmental disorders, including autism spectrum disorder (ASD) (*37*), ADHD(*38*), Tourette Syndrome(*39*), and schizophrenia (*40*). Altered striatal functional connectivity (*41*) and broad alternations in cortico-striatal white matter (*42–44*) have also been linked with repetitive behaviors and autism. However, little is known about how the putamen’s tissue microstructure and cortical connectivity develop during the first year of human life. Histological studies indicate that the broad cortex–striatum–pallidum–thalamus– cortex circuitry is established by term (*45*). However, it is unknown if the putamen’s spatial organization is already established at birth, or its microstructure and connectivity continue to develop during infancy. Here, we address this question by mapping putamen’s macro- and microstructure and cortico-putamen connectivity from birth through the first year of life and comparing these developmental trajectories with those in adults.

Regarding microstructural development, one hypothesis is that because the putamen emerges *in utero* and gets a head start, its microstructure is qualitatively and quantitatively mature at birth. This hypothesis predicts a spatial gradient along the AP axis with denser microstructure in posterior than anterior putamen and microstructural tissue density is quantitatively similar to that in adults. A second hypothesis is that the microstructure develops after birth as experience grows. In this scenario, there are two possibilities whereby (i) gradients exist at birth along the AP axis, but overall tissue is less dense in infants than adults, and tissue density increases postnally, or (ii) AP gradients are absent at birth and microstructure is less dense, with AP gradients becoming steeper and tissue becoming denser postnatally. On the question of maturation of WM connections, we make similar qualitative and quantitative predictions. The first possibility is that cortico-putamen connectivity is established *in utero*, consistent with putamen’s early emergence and hence connectivity at birth is adultlike with a topographical organization in which the anterior putamen is connected to frontal areas, and posterior putamen is connected to motor and somatosensory areas. Alternatively, cortico-putamen connectivity is immature at birth and is progressively refined postnatally, with two possible scenarios: (i) initial exuberant (*46*) widespread connectivity between cortex and putamen that is later pruned to establish adultlike topography or (ii) differential increases in cortico-putamen WM connections to sensorimotor regions as differential functional demands in sensorimotor regions to support crawling, walking, or visual recognition emerge during infancy (*47–53*).

To test these developmental hypotheses, we collected anatomical, quantitative, and diffusion MRI data in 44 infants (N_female_= 19) across 82 longitudinal sessions (age-range: 9-479 days) during infants’ natural sleep, and from 20 neurotypical adults (age-range: 19-42 years, N_female_= 12). Anatomical T_1_/T_2_ weighted images were automatically segmented into white and gray matter and subcortical structures using infant(*54*) and adult(*55*) FreeSurfer (**Supplementary Fig. 1** and **Methods**) from which we spatially delineated the putamen. We then divided each putamen into seven equidistant segments along its long length AP axis (**Fig. 1a**). We first examined the development of the volume of the putamen’s segments. Next, we examined the development of microstructural gradients using quantitative MRI metrics R_1_ and R_2_* along the AP axis. Then, we used diffusion MRI and tractography to determine the topographic organization and development of WM tracts between the putamen and cortex.

**Figure 1.**
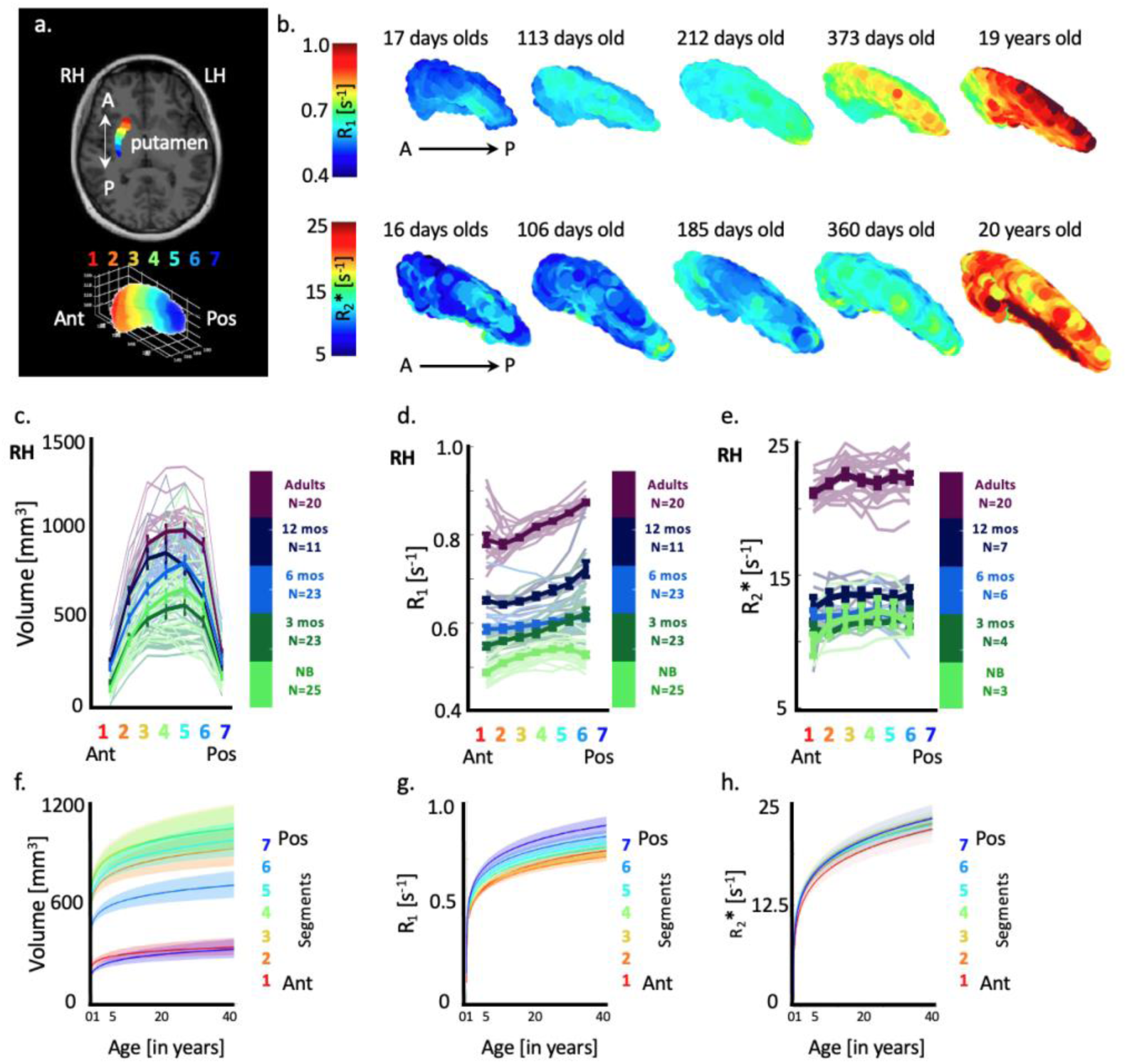
Spatial gradients in putamen’s microstructure along the anterior-posterior (AP) axis. a) Parcellation of the putamen along its AP axis into seven equidistant segments (from red to blue (A to P) for measuring gradients based on prior work(*22*). b) Three-dimensional maps of the right putamen in a sample newborn, 3-month-old (mo), 6 mo, 12 mo, and an adult) showing R_1_ (top row) and R_2_* (bottom row) along its AP axis; warmer colors represent denser microstructure. c) Volume changes per segment in newborns, 3 mo, 6 mo, 12 mo, and adults along putamen’s AP axis. Thinner lines: represent each participant’s volume trend. Error bars: standard error across participants per age group. d.) R_1_ gradients in newborns, 3 mo, 6 mo, 12 mo, and adults along putamen’s AP axis. e) same as in d for R_2_*. f) Curves representing volume changes as a function of age per putamen segment. g-h) same as in f for R_1_ and R_2_*. RH: right hemisphere. Left hemisphere (LH) data in **Supplementary Fig. 2.**

## Results

### Postnatal maturation of the human putamen follows an anterior-posterior microstructural gradient

**Figure 1b** shows R_1_ and R_2_* as 3-dimensional maps along the putamen, in example newborns, 3-month-olds, 6-month-olds, 1-year-olds and adults. The data revealed that R_1_ and R_2_* increase from infancy to adulthood, suggesting development of tissue microstructure. R_1_ also illustrates a spatial gradient along the AP axis of the putamen in all ages, with higher R_1_ values in posterior than anterior putamen (**Fig. 1b** top row). R_2_* spatial gradients along the AP axis were less prominent compared to R_1_ (**Fig. 1b** bottom row).

Next, we quantified the development of R_1_ and R_2_* along the AP axis of the putamen. As in prior work(*22*), we segmented the putamen into 7 equally-spaced segments (**Fig. 1a** and **Methods**) and estimated the volume [mm^3^], mean R_1_ and mean R_2_* [s^−1^] per segment and participant. For conciseness, we report right hemisphere data in the main manuscript and left hemisphere data in supplementary figures. We find no significant differences between hemispheres.

We find that the volume of all putamen segments increase with age (**Figs. 1c, f**) (significant main effect of age: RH: *β* = 49.3, SE = 10.7, t_710_ = 4.62, p = 4.47×10^−6^, 95% CI = [28.4 70.2]) with no significant differences in development across segments (no age by segment interaction, ps>0.05, full statistics in **Supplementary Table 1,** left hemisphere data in **Supplementary Fig. 2**), even as there are differences in volume across segments.

In all putamen segments along the AP axis, R_1_ is larger in adults (**Fig. 1d**, purple lines) than in infants (**Fig. 1d**, green-blue lines, significant main effect of age: RH: *β* = 4.23×10^−2^, SE = 1.40×10^−3^, t_710_ = 29.67, p = 1.83×10^−126^, 95% CI = [3.95 ×10^−2^ 4.5×10^−2^]), suggesting denser microstructural tissue in adults’ than infants’ putamen. Across all participants even in newborns, there is an R_1_ gradient along the AP axis where there is a progressive increase in R_1_ from the anterior to the posterior segments of the putamen (significant main effect of segment: RH: *β* = 2.8 ×10^−3^, SE = 1.40×10^−3^, t_710_ = 2.06, p = 0.039, 95% CI = [1.39×10^−4^ 5.55×10^−3^]). This gradient, however, is shallower in infants and increases from birth to one year of age to adulthood (significant interaction between age and segment: RH: *β* = 1.30×10^−3^, SE = 2.36×10^−4^, t_710_ = 5.48, p = 6.07×10^−8^, 95% CI = [8.28×10^−4^ 1.80×10^−3^]). Plotting R_1_ as function of age, per segment, revealed an anterior to posterior gradient, with steeper developmental slopes in the posterior than anterior segments (**Fig. 1g**) highlighting that microstructural growth is not uniform but varies as a function of spatial location along the AP axis of the putamen (full statistics in **Supplementary Table 2**). We also examined the development of R_1_ along equally spaced segments along the inferior-superior (IS) and medial-lateral (ML) axes. In all axes, R_1_ progressively increases from newborns to 1-year-olds and from 1-year-olds to adults (**Supplementary Fig. 3, Supplementary Table 2:** full statistics). Along the IS axis, there is no R_1_ gradient in newborns, and from infancy to adulthood an inferior-to-superior R_1_ gradient develops (**Supplementary Figs. 3a,b**). Along the ML, R_1_ is higher in the medial than lateral putamen in infants, but this gradient reverses by adulthood, with adults showing higher R_1_ in lateral than medial putamen (**Supplementary Figs. 3e,f**).

R_2_* also develops from infancy to adulthood (**Fig. 1e**, purple lines) than in infants **Fig. 1e**, green-blue lines) (significant main effect of age: RH: *β* = 2.05, SE = 0.12, t_276_ = 16.80, p = 4.03×10^−44^, 95% CI = [1.81 2.29], no significant main effect of segment or interaction between age and segment, ps>0.05, **Supplementary Table 3,** left hemisphere: **Supplementary Fig. 2**). However, R_2_* development in infancy is different than R_1_ development. R_2_* shows little development during the first six months, a minor development from six to 12 months, and a large development between infancy and adulthood and no developmental spatial gradient along the AP axis (**Fig. 1h**). We find a similar R_2_* developmental pattern along the IS axis of the putamen **(Supplementary Figs. 3c,d)** but a more graded development of R_2_* along the ML axis **(Supplementary Figs. 3g,h, Supplementary Table 3:** full statistics).

Combined, our macro and microstructural findings revealed that the putamen’s volume grows, and its tissue becomes denser from infancy to adulthood as R_1_ progressively increases. It is interesting that spatial gradients of R_1_ and R_2_* develop non uniformly and progressively across the physical axes of the putamen. These data suggest that although the putamen emerges early *in utero*, its microstructure is not mature as birth and undergoes extensive, non-uniform growth along its main axes.

### Cortico-putamen white matter tracts are topographically organized along the anterior-posterior axis in adults but are diffuse in infants

We examined the topographic organization of cortico-putamen WM tracts and their development. Using dMRI data and tractography, we generated the whole-brain, WM connectome of each participant and session, identified the tracts between each putamen segment and cortex, and quantified the distribution of endpoint densities across the entire brain, or the connectivity profile (CP) of each segment resulting in seven cortico-putamen CPs per segment/participant (**Fig. 2** and **Methods**).

**Figure 2.**
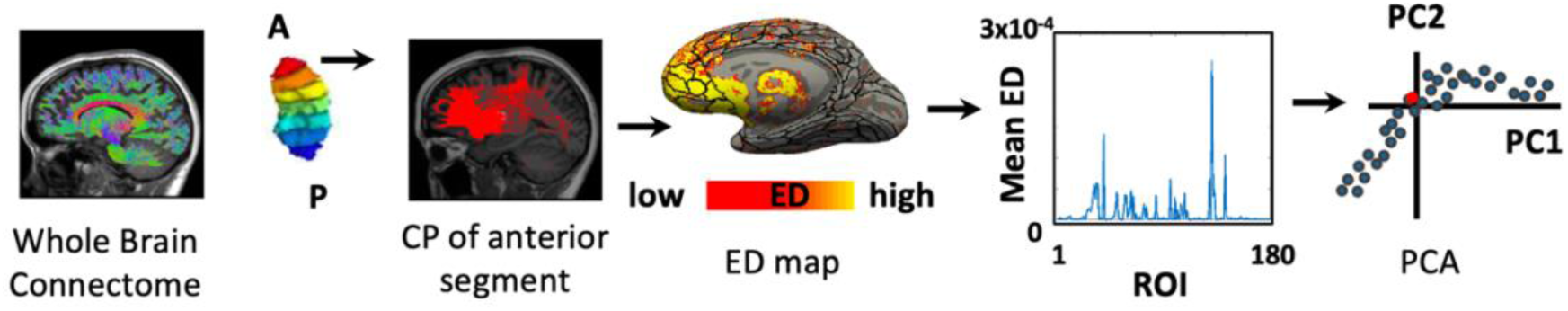
Diffusion MRI data preprocessing pipeline. Schematic showing the analysis pipeline establishing the white matter (WM) connectivity profiles (CP) for putamen segments. All analyses are done within each individual’s brain. From left to right. (1) Using dMRI and mrTrix3 and anatomically constrained tractography, we generate the participant’s whole brain connectome. (2) We intersect each segment of the putamen with the whole brain connectome and obtain the WM connections of that segment (example shows the WM connections of the most anterior segment). (3) We project the endpoints of the tracts on the cortical surface using tract density imaging (TDI) and mrTrix3 (red tracts projected on the cortex) and calculate the distribution of the endpoints across all cortical surface vertices by dividing the endpoint map by the total number of endpoints, resulting in an endpoint density (ED) map that sums to 1 (yellow represents high ED). We then used the Glasser atlas (*59*) to delineate 180 cortical regions of interest (ROI) spanning the entire cortex (black outlines on the ED surface map). (4) We calculate the mean ED per ROI, resulting in a vector of ED, which is the connectivity profile (CP) of that segment in that participant. (5) Finally, we applied principal component analysis (PCA)) on the CPs of all segments and participants, reducing the data dimensionality, highlighting the major axes of variance (e.g., PC1 vs PC2 are the two axes explaining most variance in the data) and test if CPs are separated by age and segment. Each dot in this PCA plot is a CP of one segment and participant. Red dot: CP of the anterior putamen segment for a single participant.

**Figure 3** visualizes the cortico-putamen tracts in sample infants and adults, coloring the tracts by their respective putamen segment. Results showed a clear topographic organization in adults, as anterior segments of the putamen have more tracts to frontal regions of the brain (**Fig. 3a**, top row, red-orange connections) and posterior segments have more tracts to sensorimotor-premotor regions which are more posterior (**Fig. 3a**, top row, green-blue connections). This pattern is consistent in all adults (**Supplementary Fig. 4**). At birth, this characteristic topographic, adult-like color gradient is less apparent (**Fig. 3a**, bottom row). While the anterior segments are connected to anterior brain regions (**Fig. 3a**, bottom row, red and orange tracts) as in adults, the posterior segments of the putamen in infants are also connected to the frontal cortical regions (**Fig. 3a**, bottom row, blue tracts). This makes the overall white matter topography in infants appear more diffuse than adults, and this pattern remains visible throughout the first year of life (see more example participants in **Supplementary Fig. 4**). Our observations suggest that cortico-putamen tractography is topographically organized in a clear front-to-back manner in adults, but this organization is not fully mature in infancy.

**Figure 3.**
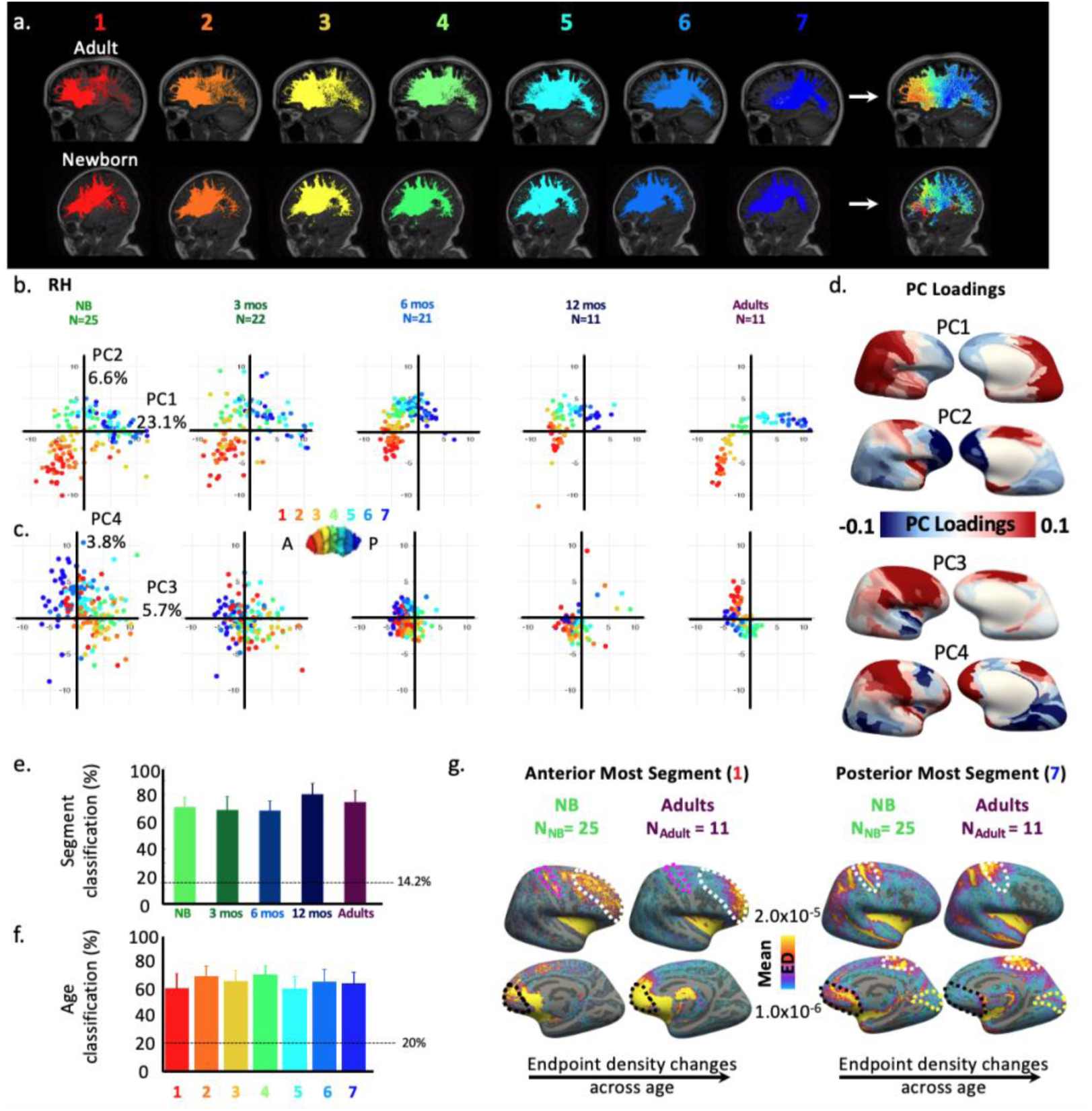
Topographical organization of cortico-putamen WM tracts along the anterior-posterior axis is more distinct in adults than infants. a) Sagittal images showing WM tracts between each putamen segment along its AP axis. Tracks are colored by segment from anterior in red to posterior in blue. Right most images show all segment tracts together. Top: sample adult (28 years old). Bottom: sample newborn (21 days old). b) Coefficients of WM connectivity profiles (CP) on the first two principal components (PC1 and PC2). Each dot: CP of one segment and one participant. Dots are colored by segment color (anterior to posterior: red to blue). Each panel shows a different age groups, light green: newborns, dark green: 3-months-old (mos), light blue: 6 mos, dark blue: 12 mos, purple: adults c) Same as in b for PC3 and PC4. d) Individual PC1-PC4 loadings for the 180 Glasser Atlas ROIs (*56*) projected on the average adult FreeSurfer surface (blues: negative loadings, reds: positive loadings). e) Classification accuracies for classifying segments from connectivity profiles. Dashed line reflects chance level. Error bar: standard error across segments. f) same as in e for classification of age. Error bar: standard error across age groups. g) Group maps of mean endpoint density (ED) of the anterior most putamen segment (left panels) and the posterior most segment (right panels) in newborns (N=25) and adults (N=11) projected on the average adult FreeSurfer surface, top row: lateral surfaces; bottom row: medial surfaces. Yellow indicates larger ED. RH: right hemisphere. Left hemisphere data in **Supplementary Fig. 5**.

We next applied a data driven PCA approach to determine the major spatial and developmental trends of the putamen WM connections. We asked if the putamen segments have separable connectivity profiles (CP), and if the CPs differ across age groups (**Fig. 2** and **Methods**). Projecting the CPs of each segment on the first two principal components (PCs, PC1 explains 23.1% of the variance, and PC2 explains 6.6% of the variance) per age group (newborn, 3-, 6-, 12-month, adults) and coloring them by segment color, reveals the following results (**Fig. 3b**): First, in all age groups, there is a clear segment-wise organization of CPs. CPs of the same segment are clustered together, are separate from CPs of other segments, with the most distinctly separable organization in adults consistent with **Figure 3a-**top row. Second, there is a spatial gradient of CPs as the CP progressively changes from the anterior to posterior putamen segment along PC1/PC2. In particular, anterior segments (1–2) have negative coefficients along PC1 and PC2, but posterior segments (5–6) have positive coefficients along PC1 and PC2. We projected each PC’s loading onto the average adult FreeSurfer brain to determine connectivity to which cortical regions may drive these differences (**Fig. 3d**). The data show that anterior putamen segments show higher connectivity to prefrontal cortex and that posterior putamen segments show higher connectivity to sensorimotor cortex along the occipital, parietal, and central/precentral sulci and gyri.

Examination of the organization of the CPs along the next two PCs, PC3 and PC4 (PC3 explains 5.7% variance and PC4 explains 3.8%) still shows clustering by segment but not in a topographic way (e.g., CP of the most anterior and posterior segments are similar along these dimensions, **Fig. 3c**). Additionally, we find age-related differences in the dispersion of CPs. Adults show a more clustered and structured organization than infants along PC3 and PC4. For example, along PC3, CPs are clustered and have largely negative coefficients in adults but are scattered with positive coefficients for segments 1-5 and negative coefficients for segments 6-7 in newborns and 3-month-olds. As the biggest difference between age groups is apparent in segments 1-5 showing negative coefficients in adults on both PC3 and 4, but positive coefficients in young infants, we examined the loadings of PC3 and PC4 on the brain. **Fig 3d** illustrates that newborns and 3-months olds have more connections between these segments to prefrontal cortex and central/precentral sulci and gyri and less connections to visual cortex in the occipital lobe than adults. Similar results were found in the left hemisphere (**Supplementary Fig. 5**).

Next, we utilized a leave-one-out classification approach to test if a classifier trained on CPs of *n*−1 participants can predict the putamen segments or participant’s age from left-out CPs. Classification used the information across 36 PCs that explained more than 80% of the variance in the data. Overall, both segments (%mean accuracy±standard deviation: RH: 72.08±2.05%, chance: 14.28%) and age (%accuracy RH: 65.86±3.0%, chance: 20%) can be classified significantly above chance (**Figs. 3e,f**, full confusion matrices in **Supplementary Fig. 6**). This highlights that the heterogeneity in putamen’s anterior versus posterior CPs contains reliable information about the spatial and developmental organization. Specifically, the putamen has a spatially organized topography, and it is developmentally reorganized. Moreover, this heterogeneity stems from differences in the ED of tracts between various brain regions across the anterior and posterior segments (**Fig. 3g**). For example, endpoint density (ED) of tracts between dorsolateral prefrontal and somatosensory cortex of the anterior most segments (**Fig. 3g**, white and magenta dotted lines) are diffuse in newborns, and this spread reduces in adults. ED of tracts between the posterior most segment and visual and sensorimotor cortex (**Fig. 3f**, yellow/white dotted lines respectively) increase from birth to adulthood, but ED between the anterior cingulate (**Fig. 3f**, black dotted lines) increases with anterior most and decreases with the posterior most segment respectively.

Overall, the tractography data highlight that the general topography of cortico-putamen connectivity is present in newborns, but it undergoes development from infancy to adulthood, and it is not adultlike at birth. In adults, the anterior putamen connects to prefrontal regions and posterior putamen to sensorimotor and temporal areas, and this distinct topographical organization is stable and replicable in adults. During infancy, connections are more diffuse as there is a higher endpoint density between the anterior putamen and prefrontal and somatosensory areas, which reduces with age, and lower endpoint density with the posterior putamen and visual cortex which increases with age, suggesting heterogeneous shifts in CPs. Critically, CPs of the putamen alone predict both the spatial location along the putamen and the participants’ age.

### Overconnectivity between putamen and dorsolateral prefrontal in ASD adults with high social impairment scores

Our finding that cortico-putamen white matter connectivity becomes more topographically organized from infancy to adulthood raises the possibility that there may be deviations from this maturation in neurodevelopmental disorders such as autism. Based on prior work (*41*, *42*, *57*), we hypothesized that adults with ASDs may show aberrant cortico-putamen connectivity profiles compared to typically developing adults. To test this hypothesis, we used an open-source diffusion imaging dataset (Autism Brain Imaging Data Exchange II: ABIDE-II, from the Barrow Neurological Institute, see **Methods**) along with its corresponding behavioral metric representing the total social responsiveness scale (SRS). Higher SRS indicate higher social impairment. This data includes total of 58 participants (age-range: 18-62 years, all participants in this dataset are males). However, for our analysis, we selected the ASD participants with SRS scores > 90 and TC participants with SRS scores < 40 to avoid any overlapping SRS scores, resulting in a data set with 20 ASD and 21 typical controls adults (**Methods**). We ran a similar analysis of WM cortical-putamen connections by putamen segment as in our developmental data and tested if CPs differed between participants with ASD and controls.

The cortico-putamen connections in the typical controls in the ABIDE-II data replicate our adult findings (**Fig. 4a**, bottom). Contrary to our hypothesis, adults with ASD (**Fig. 4a**, top row) show largely similar cortico-putamen WM connections to typical adults (**Fig. 4a**, bottom row). Adults with ASD have a clear topographic organization with anterior segments of the putamen having more tracts to frontal regions (**Fig. 4a**, top row red-orange connections) and posterior segments having more tracts to sensorimotor-premotor regions (**Fig. 4a**, top row, green-blue connections).

**Figure 4.**
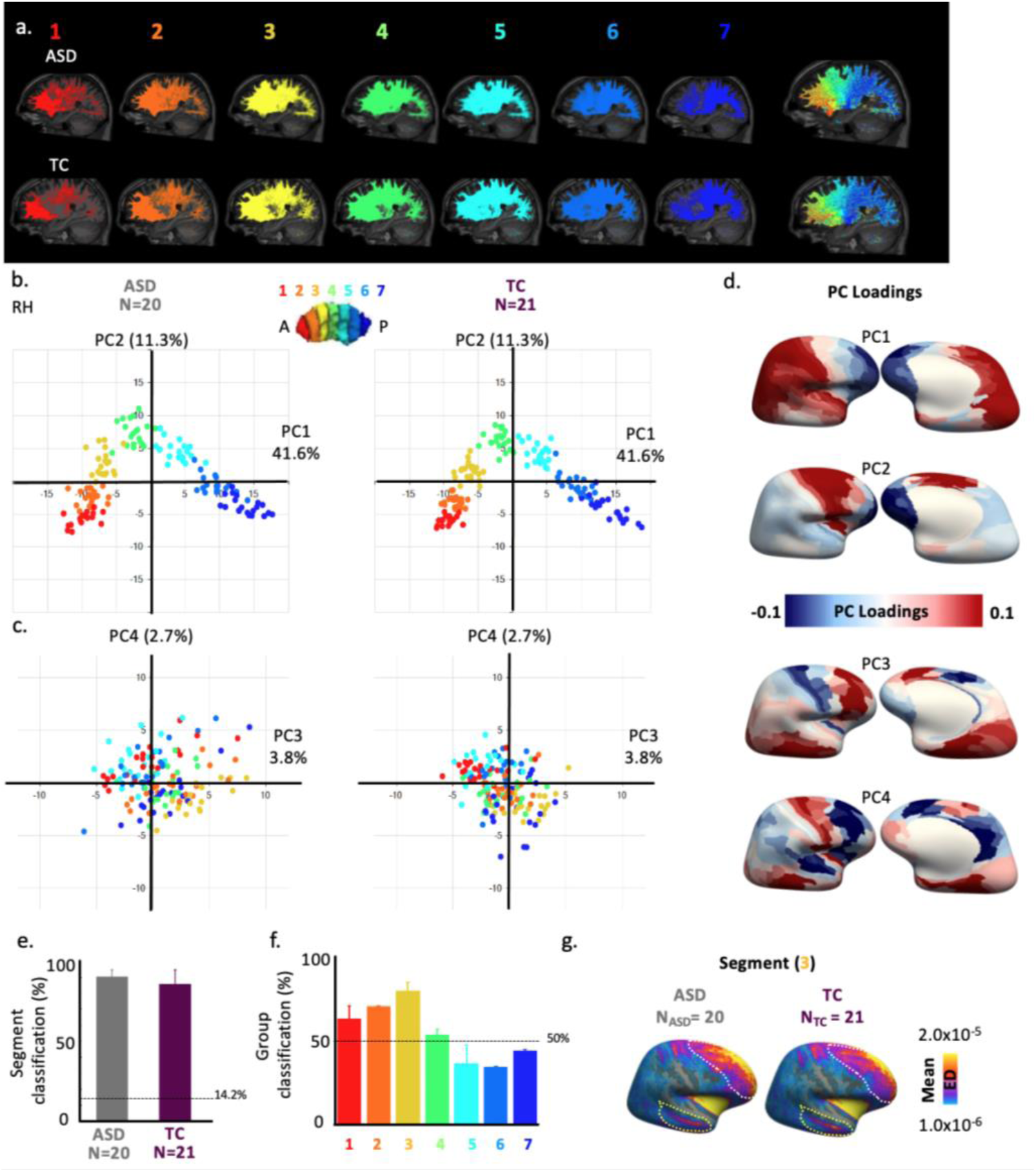
Cortico-putamen tractography in ASD adults is topographically organized like controls, with more diffuse connectivity to dorsolateral-prefrontal cortex. a) Sagittal images showing white matter fiber tracts between each putamen segmentand cortex.Top: example adult with ASD. Bottom: example typical control (TC). Tracks are colored by segment from red anterior in red and posterior in blue. Right panel shows all putamen connections together. b) Coefficients of connectivity profiles of the putamen segments projected on the first two principal components (PC1 and PC2). Left: adults with ASD (N_ASD_=20), right: typical controls (N_TC_=21). Each dot is a connectivity profile of a segment in an individual. CPs are colored by segments, red: anterior most to blue: posterior most. c) Same as in b) for PC3 and PC4. d) Individual PC1-PC4 loadings for each of the 180 Glasser Atlas ROIs projected onto the average adult FreeSurfer cortical surface. Blues: negative loadings, reds: positive loadings. e-f) Classification accuracies for classifying segments and group. g) Group maps of mean endpoint density (ED) of segment 3 along the A-P direction of the putamen in ASD and TC participants projected on average adult FreeSurfer cortical surface. Yellow indicates larger ED. Left hemisphere data in **Supplementary Fig. 7.**

PCA analysis of the ABIDE-II data shows a clear segment-wise and topographic organization of CPs visible across first two PCs (PC1 explains 41.6% and PC2 explains 11.3% variance) (**Fig. 4b**). First, in both adults with ASD and typical controls there is a clear segment-wise organization from anterior to posterior segments, as CPs of the same segment were clustered together and separate from other segments in both groups. Second, there is a spatial gradient in the clustering as anterior segments showed negative coefficients along PC1 and PC2, middle segments showed positive coefficients along PC2, and posterior segments show positive coefficients along PC1 and negative coefficients along PC2. Indeed, examining PC1 and PC2 loadings on the brain reveals that in both adults with ASD and controls, the two anterior segments of the putamen show higher connectivity to the prefrontal cortex and lower connectivity to the occipital lobe (**Fig. 4d**), but the two posterior segments show higher connectivity to the occipital and temporal lobe and lower connectivity to precentral gyri and sulci as well as prefrontal cortex (**Fig. 4d**).

To test if the next two PCs showed differences between groups, we projected the CPs on PC3 and PC4 (PC3 explains 3.8% variance and PC4 explains 2.7%) (**Fig. 4c**). Along PC3, typical control adults showed a more clustered organization than adults with ASD participants that have more scattered and positive coefficients along PC3.

PC3 loadings were positive for dorsolateral prefrontal, superior temporal areas and visual areas and were negative for motor and somatosensory cortex (left hemisphere data in **Supplementary Fig. 7**).

Using a leave-one-(participant)-out classifier, we next tested if a classifier could predict the putamen segments or participant’s group (ASD versus TC) from left-out CPs. Overall, we find that segments are classified well above chance (%mean accuracy±standard deviation: RH: 91.95±1.37%, chance: 14.28%, **Fig. 4e**). Group (ASD/TC) is only classified above chance using CPs of the three most right anterior segments (**Fig. 4f**), with the highest classification accuracy of group using the CP of segment 3 (%mean accuracy RH: 80.35±7.57%, chance: 50%). Examining ED tracts between putamen’s segment 3 and cortex in ASD versus TC revealed that ED in dorsolateral prefrontal cortex is more diffuse and widespread in adults with ASD than that in TC individuals (**Fig. 4g**).

To summarize, tractography profiles of the putamen with cortex even though appear qualitatively similar in ASD and age-matched typical control participants, there are subtle distinctions across brain areas in ASD versus controls. Specifically, in adults with ASD tracts between anterior putamen and prefrontal areas, related to executive functioning, are more diffuse and widespread than that in controls. Our results suggest that white matter tractography of putamen to different brain areas may provide clues to neurodevelopmental disorders such as autism.

## Discussion

We examined the development of the putamen’s volume, microstructure, and WM connectivity with cortex during the first year of human life and compared to its organization in adults. We find increasing volume and tissue microstructure, accompanied by a sharpening of R_1_ microstructural gradients related to tissue density along the AP axis from birth to 12 months. Additionally, R_2_*, which is modulated by iron, increases primarily between age one and adulthood, producing a later lateral–medial gradient. Cortico-putamen connectivity is diffuse in infants, with frontal cortex broadly connecting to anterior and posterior putamen but becomes clearly topographical along an AP axis in adults, with frontal regions connecting to the anterior putamen and sensorimotor cortex connecting to the posterior putamen. Adults with ASD show similar topographical organization of cortico-putamen tracts as neurotypical adults but have diffuse connectivity between the anterior putamen and prefrontal regions involved in executive function. Our findings reveal postnatal refinement of the putamen’s microstructure and connectivity during the first year of life and provide a neurotypical developmental timeline that can serve as a baseline for understanding the emergence of its function and atypical development which has implications for neurodevelopmental disorders associated with the putamen such as ASD.

Using multiparametric MR metrics, R_1_ and R_2_*, we advance the field from just measuring the putamen’s volumetric growth (*27–33*), to characterizing the developmental trajectory of the putamen’s microstructural tissue properties during the first year of life. These complementary metrics—R_1_, sensitive to myelin and dendritic arborization (*18*, *20*, *21*, *58–61*), and R_2_*, sensitive to iron concentration(*18*, *19*, *58*)—illustrate the developmental trajectory of the putamen’s microstructure during the first year of life. We find that despite the putamen’s early emergence *in utero,* its microstructure is immature at birth as R_1_ is low in newborns and does not vary much spatially. During the first year of life and into adulthood, R_1_ progressively increases and more so in the posterior than anterior segments of the putamen, generating a pronounced R_1_ gradient along the putamen’s AP axis in adults. R_2_* in the putamen follows a distinct developmental trajectory, with shallow spatial gradients at birth, little to no changes during the first year, followed by a later substantial maturation from 1-year to adulthood. This differential development leads to different spatial gradients in adulthood, where the primary gradient for R_1_ is anterior-posterior and the primary gradients for R_2_* are along the inferior-superior and lateral-medial axes. Our findings suggest that while the putamen is not microstructurally homogeneous at birth, its microstructural growth, which may involve increases in myelin and dendritic and axonal arborization, follows a different trajectory from iron accumulation. These differential developments may support differential functional and metabolic demands during infancy.

Our second developmental finding is that WM cortico-putamen connections are also immature at birth. While different segments of the putamen along the AP axis have different white matter connections in both newborns and adults, the topographic organization of the cortico-putamen connections is more distinct in adults than infants. In newborns, cortico-putamen connections are more diffuse than adults, as newborns have higher endpoint density than adults between anterior putamen and prefrontal cortex and posterior putamen and the anterior cingulate but lower endpoint density than adults between posterior putamen and somatosensory and visual cortices. While there is refinement of cortico-putamen connections over the first year of life, the topographic organization of cortico-putamen connections remains immature at 1-year of age. The spatially diffuse WM connectivity during infancy that becomes more topographically organized over development is consistent with the exuberance hypothesis (*46*, *62*) which proposes that early brain development is characterized by an overproduction of connections that are subsequently refined over development. However, as dMRI data cannot reveal the cellular mechanisms of WM refinement, other cellular mechanisms may contribute to these observations including: WM myelination (*63–66*), integration of existing fibers (*63*, *66–71*), activity-dependent axonal and dendritic remodeling (*72–75*), and changes in fiber organization or geometry (*76*, *77*). Future work combining diffusion and quantitative MRI with histology are necessary for identifying the cellular mechanisms that refine the organization of cortico-putamen WM tracts.

Combined, our findings extend the prior developmental framework proposed by Kostović et al. (*45*) which suggests that the basic cortex–striatum–pallidum–thalamus–cortex circuitry is established by term, but continues to develop postnatally. Although the newborn putamen is connected to cortex at birth, its microstructural organization and connectivity are immature and refinements occur during infancy and continue throughout childhood into adulthood (*27*). We propose a novel hypothesis that the development of the putamen’s microstructure and its WM connectivity with cortex may support the growing functional demands during infancy. As infants acquire complex somatosensory, and visual-motor skills (*52*, *53*, *78*, *79*), cortical connections from these regions to the posterior putamen may increase alongside the microstructural growth of the posterior putamen during the first year of life. Refinement of connections from frontal regions to the anterior putamen regions and slower microstructural growth of the anterior putamen may be tied to the prolonged maturation of higher-order cognitive functions that the putamen may support. Future studies combining functional and quantitative MRI in children and adolescents will be important for testing these structure–function relationships between the putamen, other basal ganglia nuclei, and cortex.

Comparing adults with ASD to neurotypical adults revealed that adults with ASD show, like neurotypical adults, a topographic cortico-putamen WM organization along the AP axis. Nonetheless, we also observed differences in WM connections, whereby adults with ASD show higher endpoint density than neurotypical adults between the putamen’s third most anterior segment and prefrontal areas that are involved in higher-cognition and executive functions. This suggests that ASD may involve a specific pattern of aberrant cortico-putamen WM connectivity. Future research in children with ASD vs neurotypical controls can determine if this atypicality is due to developmental exuberance (*80*) or atypical pruning (*41*, *44*, *81*).

Our classification analyses suggest that cortico-putamen WM connections contain information about spatial gradients, developmental age, and neurodevelopmental differences. While the clear separability of anterior and posterior connectivity profiles in adults supports their distinct roles in cognitive and sensorimotor functions and validates our qualitative observations, the ability to classify infant age highlights that the WM connectivity has a clear developmental trajectory. Thus cortico-putamen WM connectivity patterns contain systematic information about maturational stages. In other words, an infant whose cortico-putamen WM connectivity looks substantially younger or older than expected could be identified as having atypical development. This is an important discovery as it demonstrates that cortico-putamen connectivity contains a measurable signature of brain maturation. Above-chance classification of ASD versus neurotypical adults from cortico-putamen connectivity patterns has clinical implications as they have the potential to be used as a diagnostic to identify atypical neurodevelopment. Finally, our multimodal framework using measurements of R_1_, R_2_*, and WM connections, provide a quantitative framework to evaluate the putamen not only during infancy but also in (i) childhood and adolescent development (*82–84*), (ii) individuals with neurodevelopmental disorders associated with aberrant WM connectivity including cerebral palsy(*85*), Tourette syndrome (*39*, *86*), ADHD(*38*), psychosis (*87*), Schizophrenia(*40*), obsessive-compulsive disorder(*88*), and (iii) aging disorders such as Parkinson’s disorder(*22*).

Together, our findings establish a non-invasive framework for quantifying developmental trajectories of the microstructure and cortico-striatal connectivity and provide a foundation to understand how the putamen may become functionally and spatially specialized across development. These data also provide an important foundation for understanding how early-life factors and neurodevelopmental disorders may affect the microstructural growth and connectivity of the putamen across the lifespan.

## Methods

### Participants

#### Infants and Adults

Eighty-two full-term and healthy infants (N_female_ = 35) were recruited to participate in the study. Forty-four infants (N_female_=19) provided usable data. Infants were scanned at newborn, 3 months, 6 months, and 12 months of age (age-ranges=9-479 days, Mean age±Std: 143.48±118.43 days), we obtained imaging sessions a total of 82 sessions including 25 infants who were scanned longitudinally. We excluded data from sessions when infants could not fall asleep inside the MRI scanner, which led to excessive motion and noisy data (see additional exclusion criteria in section titled “Expectant parent and infant screening procedure”). We also tested 20 adults (N_female_=12, age-range=19-42 years, Mean age±Std: 27.07±6.41 years). Adult subjects are Stanford University affiliates. All adult participants had normal or corrected-to-normal vision and provided written, informed consent. Protocols were approved by the Stanford Internal Review Board on Human Subjects. The participant population was racially and ethnically diverse, reflecting the population of the San Francisco Bay Area, Hispanic, Asian Caucasian, and multiracial participants.

### Expectant parent and infant screening procedure

Expectant parents and their infants in our study were recruited from the San Francisco Bay Area using social media platforms. We performed a two-step screening process. First, parents were screened over the phone for eligibility based on exclusionary criteria designed to recruit a sample of typically developing infants. Second, eligible expectant mothers were screened once again after giving birth. Exclusionary criteria were as follows: recreational drug use during pregnancy, significant alcohol use during pregnancy (more than three instances of alcohol consumption per trimester; more than 1 drink per occasion), lifetime diagnosis of autism spectrum disorder or a disorder involving psychosis or mania, taking prescription medications for any of these disorders during pregnancy, insufficient written and spoken English ability to understand the instructions of the study, or learning disabilities that would preclude participation. Infants were excluded if they were born prior to 36 gestational weeks, with low birthweight (<5 lbs 8 oz). Additional infant exclusion criteria included the presence of congenital, genetic, or neurological disorders, visual impairments, complications at birth requiring intensive care (e.g., NICU admission), prior head injuries, or any contraindications for MRI (e.g., metal implants). Participants were compensated with 25 dollars per hour for their participation.

### Data Acquisition

All infants and adults participated in multiple scanning sessions t to obtain anatomical and quantitative MRI data. Scanning was done in a 3T GE scanner at the Center for Cognitive and Neurobiological Imaging in Stanford University. For all adult participants, we used the Nova 32-ch head coil at the CNI at Stanford University. The infant scans were collected using a custom 32-channel infant head coil developed specifically to fit newborn to one year old infants (*89*). As infants have low weight, all imaging was done with first level SAR to ensure their safety. Scanning sessions were scheduled in the evenings around the infants’ typical bedtime. Each session lasted between 2.5 h and 5 h including time to prepare the infant and waiting time for them to fall asleep. Upon arrival, caregivers provided written, informed consent for themselves and their infant to participate in the study. Before entering the MRI suite, both the caregiver and infant were checked to ensure that they were metal-free, and caregivers changed the infant into MR-safe cotton onesies and footed pants provided by the researchers. The infant was wrapped with a blanket with their hands to their sides to avoid their hands creating a loop and the researchers inserted soft wax earplugs into the infant’s ears. During sessions involving newborn infants, an MR-safe plastic immobilizer (MedVac, www.supertechx-ray.com) was used to stabilize the infant and their head position. Once the infant was ready for scanning, the caregiver and infant entered the MR suite. The caregiver was instructed to follow their child’s regular sleep routine. When the infant was asleep, the caregiver placed the infant on the scanner bed. Weighted bags were placed at the edges of the bed to prevent any side-to-side movement. Additional pads were also placed around the infant’s head and body to stabilize head position. MRI compatible neonatal noise attenuators and headphones were placed on the infant’s ears, to lower sound transmission. An experimenter stayed inside the MR suite with the infant during the entire scan. For additional monitoring of the infant’s safety and tracking of the infant’s head motion, an infrared camera was affixed to the head coil and positioned for viewing the infant’s face in the scanner. The researcher operating the scanner monitored the infant via the camera feed, which allowed for the scan to be stopped immediately if the infant showed signs of waking or distress. This setup also allowed tracking the infant’s motion; scans were stopped and repeated if there was excessive head motion. To ensure scan data quality, in addition to real-time monitoring of the infant’s motion via an infrared camera, MR brain image quality was also assessed immediately after acquisition of each sequence and sequences were repeated if necessary.

### Data acquisition parameters and preprocessing

#### Anatomical MRI

T1-weighted and T2-weighted images were acquired and used for tissue segmentation. T1-weighted image acquisition used GE’s BRAVO sequence with TE=2.7ms, TR=6.7ms, echo train length = 1; voxel size = 1mm^3^; Scan time: ∼3 min. T2-weighted image acquisition used GE’s CUBE sequence with TE=122ms, TR = 3650 ms; echo train length = 120; voxel size = 1 mm^3^ ; FOV = 20.5 cm; Scan time: ∼4 min. T1-weighted and T2-weighted images were used for segmentation of gray-white matter to generate cortical surface reconstructions and delineating the putamen. All data were kept in native brain space as all analyses were performed within-subject and within-timepoint.

#### Quantitative MRI (longitudinal relaxation rate R_1_)

In all 82 out of the 82 infant sessions and all adults we obtained quantitative longitudinal relaxation rate (R_1_) which is used an *in vivo* proxy for myelin in white matter(*18*, *59*) and cortex(*20*). An inversion-recovery EPI (IR-EPI) sequence(*90*) with multiple inversions times (TI) was used to estimate quantitative relaxation time R_1_ (R_1_ = 1/T_1_) in each voxel. The IR-EPI used a slice-shuffling technique to acquire 20 TIs with the first TI = 50 ms and TI interval = 150 ms. A second IR-EPI with reverse-phase encoding direction was also acquired. Other acquisition parameters were voxel size = 2 mm^3^ isotropic; number of slices = 60; FOV = 20 cm; in-plane/through-plane acceleration = 1/3; Scan time: 1 min and 45 sec. To obtain R_1_ maps, we first performed susceptibility-induced distortion correction on the IR-EPI images using FSL’s top-up(*91*) and the IR-EPI acquisition with reverse-phase encoding direction. We then used the distortion corrected images to fit the T_1_ relaxation signal model using a multi-dimensional Levenberg-Marquardt algorithm. In an inversion-recovery sequence, the signal *S(t)* has an exponential decay over time (*t*) with a decay constant *T_1_*

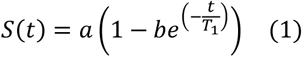

In Eq 1, *a* is a constant that is proportional to the initial magnetization of the voxel and *b* is the effective inversion coefficient of the voxel (for perfect inversion *b* = 2). We applied an absolute value operation on both sides of the equation and used the resulting equation as the fitting model because the magnitude images were used to fit the model. The magnitude images only keep the information about the strength of the signal but not the phase or the sign of the signal. The output of the algorithm is the estimated quantitative T_1_ value per voxel. From the T_1_ estimate, we calculate R_1_ (R_1_ = 1/T_1_) as R_1_ is directly proportional to macromolecular tissue volume. QMRI data fitting is performed using the brain mask and not the whole head.

Since we collected the qMRI data around the infant’s typical bedtime, and only when the infant was asleep inside the scanner, we did not perform any additional motion correction on the qMRI scan as all scans were collected with sleeping infants. However, if we noticed any ringing in and around the scans due to any motion, we excluded the infant’s data from further analysis. Similar criterion was used for quality control of adult data.

#### Quantitative MRI (transverse relaxation rate R_2_*)

In a subset of our infant and all adult sessions (N_infant-session_=20 and N_adult_ =20), we also obtained a 3D Fast Gradient Echo with Multi-Echo sequence to acquire the data for quantitative R_2_* measurement. Acquisition parameters are as follows: FOV = 20cm, phase FOV = 16cm, acquisition matrix = 200×160, number of slices = 116, slice thickness = 1mm, flip angle = 15deg, pixel bandwidth = 488Hz/pixel, in-plane acceleration = 2, number of echoes = 12, min TE = 2.9ms, delta TE = 3.1ms, TR = 46ms. Real and imaginary DICOM images for each individual echo are reconstructed. We then used the MEDI Toolbox and its ARLO algorithm for fitting the R_2_*(*92*).

#### Diffusion MRI (dMRI) acquisition

In 79 out of 82 infant sessions and 11 out of 20 adult sessions, we obtained dMRI data with the following parameters: multi-shell, #diffusion directions/b-value = 9/0, 30/700, 64/2000; TE = 75.7 ms; TR = 2800 ms; voxel size = 2 mm^3^ isotropic; number of slices = 60; FOV = 20 cm; in-plane/through-plane acceleration = 1/3; scan time: 5:08 min. We also acquired a short dMRI scan with reverse phase encoding direction and only 6 b = 0 images (scan time 0:20 min).

#### Generation of cortical surfaces

##### For infants

We generated gray and white matter tissue segmentations using the T1- and T2-weighted images. Multiple steps were applied to generate an accurate segmentation of each infant’s brain at each timepoint (**Supplementary Fig. 1**): (1) An initial segmentation of gray and white matter was generated from the T1-weighted brain volume using infant FreeSurfer’s automatic segmentation code ((infant-recon-all; version freesurfer-linux-centos7_x86_64-infant-dev-4a14499-20210109; https://surfer.nmr.mgh.harvard.edu/fswiki/infantFS(54)). (2) A second segmentation was done using both the T1- and T2-weighted anatomical images and the brain extraction toolbox (Brain Extraction and Analysis Toolbox, iBEAT, v-2.0 cloud processing, https://ibeat.wildapricot.org/). iBEAT V2.0 was specifically designed for processing infant brain MRI data using contrasts properties of both T1w and T2w images, and employs deep learning techniques trained on infant data to handle the unique challenges of low contrast between gray and white matter in developing brains. Through visual inspection, we determined that iBEAT provided a more accurate segmentation of the gray-white matter boundaries in our low-contrast infant images compared to the infant FreeSurfer approach. (3) The iBEAT segmentation was further manually corrected to fix segmentation errors in the white and gray matter (such as mislabeled white matter voxels or holes) using ITK-SNAP (http://www.itksnap.org/). (4) The iBEAT corrected segmentation was reinstalled into FreeSurfer. The reinstalling process allowed us to obtain clean surface maps, and more accurate macroanatomical features required for our analysis. The resulting segmentation in FreeSurfer format was used for analyses.

##### For adults

We used T1-weighted anatomical scan and FreeSurfer’s (version 7.2.0) automatic segmentation code (https://surfer.nmr.mgh.harvard.edu/) to segment gray and white matter surfaces per adult brain. Gray matter segmentations were checked to fix segmentation errors using ITK-SNAP (http://www.itksnap.org/) and corrected segmentations were reinstalled into FreeSurfer.

#### Segmentation of the putamen

In each participant and session, the putamen was isolated using infant/adult FreeSurfer’s automatic segmentation code. The individual putamen was further manually corrected to fix segmentation errors and mislabeled voxels using FreeSurfer’s voxel-edit tool (https://surfer.nmr.mgh.harvard.edu/fswiki/FreeviewGuide/FreeviewTools/VoxelEdit).

#### QMRI gradients within the putamen

After delineating the putamen, we estimated the microstructural gradients for R_1_ in each participant’s putamen using the automatic procedure developed in prior work (https://github.com/MezerLab/mrGrad)(22) to generate qMRI functions along the main axes of a subcortical structure per participants. Specifically, the algorithm computes the single value decomposition of the putamen’s voxel using its 3D image coordinates to find the main three orthogonal axes (i.e., the eigenvectors). The longest axis of the putamen is identified as the anterior-posterior axis, followed by the inferior-superior and medial-lateral axes. Next, per axis, we segmented the putamen with equal spacing by defining the edges as the two hyperplanes defined by the two extreme data points of the putamen with respect to the axis and by the axis as a normal to the plane. The putamen was then segmented by *n* − 1 parallel hyperplanes equally spaced between the two edges. Voxels were classified to *n* segments based on criteria of distance from planes. For our analysis, we chose *n* = 7. Per axis, segment, and participant, we measured mean R1 / R_2_*. This yielded three map-based functions of spatial position (“gradients”) along the 3 axes of the putamen.

#### Analysis of development in microstructural gradients

To quantify developmental effects, we used linear mixed models (LMMs(*93*)) as they allow explicit modeling of both within-subject effects (e.g., longitudinal measurements) and between-subject effects (e.g., cross-sectional data) with unequal number of points per participants, as well as examine main and interactive effects of both continuous (e.g., age in days) and categorical (e.g., segments) variables. To test if there were age-related differences in R_1_ and R_2_* per segment and hemisphere, we fit an LMM relating the mean metric as a function of log of age [in days] and segment using the Wilkinson notation(*94*) specify an LMM with the “fitlme” function in MATLAB (version 2020b, MathWorks, Inc.) as follows:

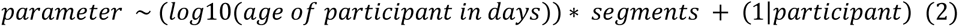

Parameters are the putamen volume, R_1_, R_2_*, which are the dependent variables, age of infant is continuous predictor [(age in days)], segment is a categorical variable (1 to 7), and the term: 1|*participant* indicates a random intercept per participant.

#### Diffusion MRI processing

Data were preprocessed using MRtrix3(*95*) (https://github.com/MRtrix3/mrtrix3) and in accordance with prior work from our lab(*65*, *66*). Data were denoised using principal component analysis(*96*). We used FSL’s top-up tool (https://fsl.fmrib.ox.ac.uk/) and one image with reverse phase-encoding to correct for susceptibility-induced distortions. We used FSL’s eddy tool to perform eddy current and motion correction, where outlier slices were detected and replaced(*97*). Finally we performed bias correction using ANTs (https://picsl.upenn.edu/software/ants/)(98). The preprocessed dMRI data were aligned to the T2-weighted anatomy for infants, and T1-weighted anatomy for adults using whole-brain rigid body registration. Alignment was checked manually for all images.

#### Generating White Matter Connectomes

We generated a whole brain white matter connectome in each session using MRTrix3. Voxel-wise fiber orientation distributions (FODs) were calculated using constrained spherical deconvolution (CSD). We used the Dhollander algorithm(*99*) to estimate the three-tissue response function. We computed FODs separately for the white matter and CSF. The gray matter was not modeled separately, as white and gray matter do not have sufficiently distinct b-value dependencies to allow for a clean separation of the signals. Finally, we generated a whole brain white matter connectome per participant’s session. Tractography was optimized using the gray/white matter segmentation from anatomical MRI data (Anatomically Constrained Tractography; ACT(*100*)). For each connectome, we used probabilistic fiber tracking with the following parameters: IFOD2 algorithm, step size of 0.2 mm, minimum length of 4 mm, maximum length of 200 mm, and maximum angle of 15°. Each connectome consisted of 5 million streamlines. Streamlines were randomly seeded on this gray-matter/white-matter interface, which is critical for accurately identifying the connections that reach regions located in the gray matter(*101*).

#### dMRI Quality Assurance

To evaluate the quality of the diffusion data we first measured the number of outliers (dMRI volumes with signal dropout measured by FSL’s eddy tool). The exclusion criterion was >5% outlier volumes. Next, we visualized in mrView the fractional anisotropy (FA) colored by direction of the diffusion tensors to validate the expected maps. That is, the existence of between hemispheric connections through the corpus callosum, and the inferior-superior directionality of the cerebral spinal tract.

#### Identifying white matter connections per putamen segment

To identify the white matter connections per putamen, we intersected the whole brain connectome generated from dMRI and tractography in each participant. We used an open source software package, FSuB Extractor(*102*) (https://github.com/smeisler/fsub_extractor) and built in MRtrix3 functions to intersect each putamen with the whole brain connectome of each participant and identify the white matter connections. In brief, the software takes in region of interest (ROI) (e.g. a single putamen segment) in the native space of each participant and projects them along the surface normal into the gray-matter-white-matter interface. It then restricts the ROI to the gray-matter-white-matter interface and then selects all streamlines that intersect with the ROI.

#### Defining endpoint connectivity profiles and principal component analysis

Quantifying the fiber tracts themselves is challenging as we observe a significant amount of overlap between the tracts originating from different anterior-posterior locations along the putamen. Hence, for each putamen segment, we quantified how it connects to the rest of the brain. Specifically, using the whole brain connectome and tract density imaging (TDI) with MRtrix3, we extracted the white matter connections of each putamen’s segment onto each subject’s cortical surface and calculated the distribution of these white matter endpoints across all cortical surface vertices. We transformed the TDI output into a distribution of endpoints by dividing the endpoint map by the total number of endpoints. This results in an endpoint density (ED) map that sums to 1 for each segment and participant. Next, we used the Glasser atlas(*56*) to delineate 180 cortical regions of interest (ROI) and obtained mean ED per ROI, resulting in a data of connectivity profiles (CP): (segment (N=7) by ROI (N=180) by participant (N=90: N_infant-session_=79; N_adult_=11). Finally, we used a data-driven approach, principal component analysis (PCA), to examine if there are segment- and age-related differences in the connectivity profiles.

#### Classification

We used an *n*-way classifier to predict the putamen segment (N=7) and age (N=5, newborns, 3-, 6-12-month-olds, and adults) of a held-out connectivity profile. We used a multiclass model for support vector machines with leave-one-(participant)-out cross-validation on all data, excluding data from the held-out participant. We then calculated classification accuracy by comparing the predicted classification to the ground truth.

#### Autism spectrum disorder (ASD) participants and Typical Controls (TC)

ASD data and the corresponding age-matched typical controls were obtained from the Autism Brain Imaging Data Exchange II (ABIDE-II, specifically data collected from Barrow Neurological Institute). All patients and controls underwent the similar protocols, including structural and diffusion imaging. Additionally, as a part of the clinical evaluations, the ABIDE-II dataset also provides behavioral measures per participant, these measures are listed here: (https://fcon_1000.projects.nitrc.org/indi/abide/ABIDEII_Data_Legend.pdf). For our study, we used a measure of social impairment as represented by the total Social Responsiveness Scale (SRS-2) which is a questionnaire that measures the presence and severity of social impairment associated with autism spectrum disorder. Specifically, it accesses social awareness, cognition, communication, motivation, and repetitive behaviors and ranges between 0 to 160, with overlaps in SRS between some typical and atypical individuals. All information on the study can be found here: https://fcon_1000.projects.nitrc.org/indi/abide/abide_II.html. Of the total of 58 participants (age-range: 18-62 years, all participants in this dataset are males) in this data set, we only selected the ASD participants with SRS scores > 90 and TC participants with SRS scores < 40 to avoid any overlapping participant scores, resulting in a data set with 20 ASD and 21 typical controls adults.

#### Acquisition parameters

All ASD and TC participants were scanned on a 3.0 Tesla Ingenia scanner, software version 5.1.9, 15-channel head coil. All T1w scan parameters are listed here: https://fcon_1000.projects.nitrc.org/indi/abide/scan_params/ABIDEII-BNI_1/anat.txt and all diffusion imaging parameters are listed here: https://fcon_1000.projects.nitrc.org/indi/abide/scan_params/ABIDEII-BNI_1/dti.txt.

#### MRI (T1w and diffusion) data processing

For the ASD data set, we used the same processing pipeline as in our data set to define the putamen in both hemispheres, divide the putamen into 7 segments, identify white matter connections per segment, generate connectomes, and define connectivity profiles.

#### Defining connectivity profiles and PCA analysis on ASD data

Similar to our data set, we obtained endpoint density maps and connectivity profiles per participant and conducted a PCA on a data of connectivity profiles to examine if there are segment- and group-related differences in the putamen-cortical connectivity profile and an *n*-way leave-one-out classifier to predict the putamen segment (N=7) and group (N=2, ASD, TC) in the ASD data set, of a held-out connectivity profile.

## Acknowledgements

This research was supported by the Stanford Wu Tsai Neurosciences Institute Big Idea and Accelerator Grants, NIH R01 EY022318 (Grill-Spector), NIH R01 EY033835 (Grill-Spector), NIH R21 EY030588 (Grill-Spector), NIH R01 HD114719 (Liao), and NIH R01 MH116173 (Setsompop). Thank you to Ahmad Allen, Karla Perez, Keithan Ducre, and Danya Ortiz for manual segmentations of the infant brains.

## Contributions

V.S.N: participant recruitment, data acquisition and preprocessing, data analysis, manuscript writing; C.T, X.Y, S.T: participant recruitment, data acquisition and preprocessing; S.K.A, E.K: diffusion data processing; H.W, N.W, C.L, X.C, K.S: MRI sequence development; A.M, E.D: discussions and code sharing on putamen sectioning; and K.G.S: overseeing all aspects of research. All co-authors read and approved the submitted manuscript.

## Competing interests

The authors declare no competing interests.

## Data availability

Source data used in the analyses and to reproduce figures have been made freely available in Github under accession code: https://github.com/VPNL/putamen_microstructuralgradients. Requests for further information or raw data should be directed to the corresponding author, Vaidehi S. Natu.

---

**Supplementary Table 1.**
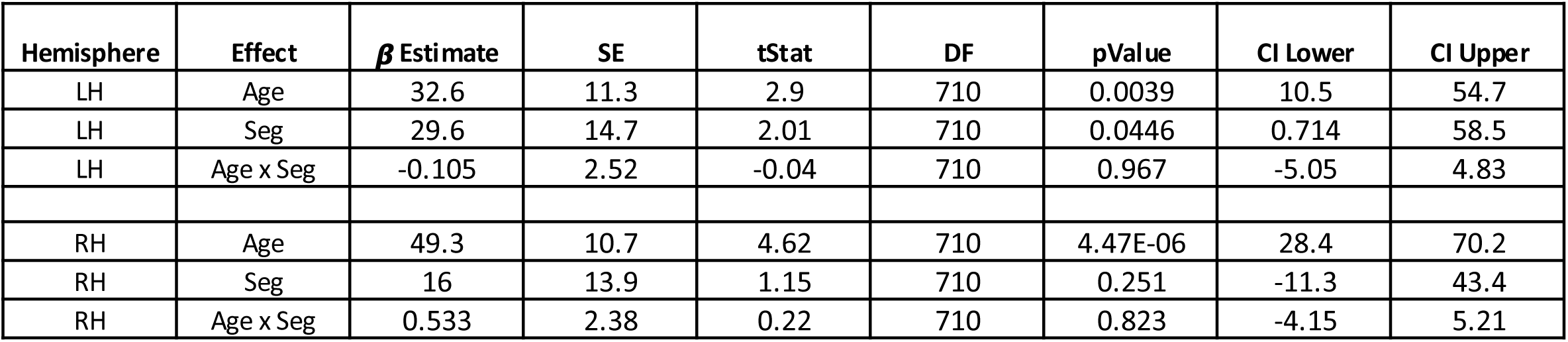
Statistical significance of the linear mixed model (LMM) quantifying the relationship between volume and age across putamen segments. LMMs quantifying, the relationship between volume and age, across the 7 segments: Volume ∼ log10(age in days) x Segment + (1|participant). Three effects are shown per model, hemisphere: main effects of age and segment and interaction between age and segment. Statistics: SE: Standard Error; DF: degrees of freedom; tstat: t-statistics; CI=confidence intervals. LH/RH: left/right hemisphere.

**Supplementary Table 2.** Statistical significance of linear mixed models (LMMs) quantifying the relationship between R_1_ and age and putamen segments. LMMs quantifying, per putamen axis (anterior-posterior: AP, medial-lateral: ML, IS: inferior-superior) the relationship between R_1_ and age, across the 7 segments. R_1_ ∼ log10(age in days) x Segment + (1|participant). Three effects are shown per model, hemisphere, and axis: main effects of age and segment and interaction between age and segment. Statistics: SE: Standard Error; DF: degrees of freedom; tstat: t-statistics; CI=confidence intervals. LH/RH: left/right hemisphere.

| Axis | Hemisphere | Effect | $\beta$ Estimate | SE | tStat | DF | pValue | CI Lower | CI Upper |
| --- | --- | --- | --- | --- | --- | --- | --- | --- | --- |
| AP | LH | Age | 4.25E-02 | 1.30E-03 | 33.76 | 710 | 8.96E-150 | 4.00E-02 | 4.50E-02 |
|  | LH | Seg | -5.00E-03 | 1.30E-03 | -3.95 | 710 | 8.55E-05 | -7.50E-03 | -2.50E-03 |
|  | LH | Age x Seg | 2.20E-03 | 2.17E-04 | 10.13 | 710 | 1.30E-22 | 1.80E-03 | 2.60E-03 |
| IS | LH | Age | 4.31E-02 | 1.30E-03 | 33.46 | 710 | 4.55E-148 | 4.06E-02 | 4.56E-02 |
|  | LH | Seg | -7.50E-03 | 1.30E-03 | -5.86 | 710 | 7.13E-09 | -1.00E-02 | -5.00E-03 |
|  | LH | Age x Seg | 2.10E-03 | 2.19E-04 | 9.45 | 710 | 4.84E-20 | 1.60E-03 | 2.50E-03 |
| ML | LH | Age | 3.44E-02 | 1.40E-03 | 25.53 | 710 | 2.02E-102 | 3.18E-02 | 3.71E-02 |
|  | LH | Seg | -2.34E-02 | 1.40E-03 | -17.21 | 710 | 1.02E-55 | -2.61E-02 | -2.07E-02 |
|  | LH | Age x Seg | 3.50E-03 | 2.34E-04 | 14.74 | 710 | 4.17E-43 | 3.00E-03 | 3.90E-03 |
| AP | RH | Age | 4.23E-02 | 1.40E-03 | 29.67 | 710 | 1.83E-126 | 3.95E-02 | 4.50E-02 |
|  | RH | Seg | 2.80E-03 | 1.40E-03 | 2.06 | 710 | 3.94E-02 | 1.39E-04 | 5.60E-03 |
|  | RH | Age x Seg | 1.30E-03 | 2.36E-04 | 5.48 | 710 | 6.07E-08 | 8.28E-04 | 1.80E-03 |
| IS | RH | Age | 3.84E-02 | 1.40E-03 | 27.56 | 710 | 2.79E-114 | 3.56E-02 | 4.11E-02 |
|  | RH | Seg | -7.90E-03 | 1.30E-03 | -5.94 | 710 | 4.51E-09 | -1.05E-02 | -5.30E-03 |
|  | RH | Age x Seg | 2.30E-03 | 2.28E-04 | 10.1 | 710 | 1.64E-22 | 1.90E-03 | 2.80E-03 |
| ML | RH | Age | 3.83E-02 | 1.50E-03 | 25.58 | 710 | 8.24E-103 | 3.54E-02 | 4.12E-02 |
|  | RH | Seg | -1.59E-02 | 1.50E-03 | -10.89 | 710 | 1.15E-25 | -1.88E-02 | -1.30E-02 |
|  | RH | Age x Seg | 1.90E-03 | 2.50E-04 | 7.41 | 710 | 3.60E-13 | 1.40E-03 | 2.30E-03 |

**Supplementary Table 3.** Statistical significance of linear mixed models (LMMs) quantifying the relationship between R2* and age and putamen segments. LMMs quantifying, per putamen axis (anterior-posterior: AP, medial-lateral: ML, IS: inferior-superior) the relationship between R_2_* and age, across the 7 segments. R_2_* ∼ log10(age in days) x Segment + (1|participant). Three effects are shown per model, hemisphere, and axis: main effects of age and segment and interaction between age and segment. Statistics: SE: Standard Error; DF: degrees of freedom; tstat: t-statistics; CI=confidence intervals. LH/RH: left/right hemisphere.

| Axis | Hemisphere | Effect | $\beta$ Estimate | SE | tStat | DF | pValue | CI Lower | CI Upper |
| --- | --- | --- | --- | --- | --- | --- | --- | --- | --- |
| AP | LH | Age | 2.38E+00 | 1.04E-01 | 22.85 | 276 | 1.32E-65 | 2.17E+00 | 2.58E+00 |
|  | LH | Seg | 1.78E-01 | 1.16E-01 | 1.53 | 276 | 1.27E-01 | -5.08E-02 | 4.08E-01 |
|  | LH | Age x Seg | -1.59E-02 | 1.56E-02 | -1.01 | 276 | 3.11E-01 | -4.66E-02 | 1.49E-02 |
| IS | LH | Age | 2.03E+00 | 1.07E-01 | 18.9 | 276 | 1.07E-51 | 1.82E+00 | 2.24E+00 |
|  | LH | Seg | -2.73E-01 | 1.33E-01 | -2.05 | 276 | 4.17E-02 | -5.35E-01 | -1.04E-02 |
|  | LH | Age x Seg | 5.81E-02 | 1.79E-02 | 3.25 | 276 | 1.30E-03 | 2.29E-02 | 9.33E-02 |
| ML | LH | Age | 2.66E+00 | 1.07E-01 | 24.96 | 276 | 9.40E-73 | 2.45E+00 | 2.87E+00 |
|  | LH | Seg | 2.76E-01 | 1.78E-01 | 1.55 | 276 | 1.21E-01 | -7.36E-02 | 6.26E-01 |
|  | LH | Age x Seg | -9.90E-02 | 2.39E-02 | -4.15 | 276 | 4.45E-05 | -1.46E-01 | -5.20E-02 |
| AP | RH | Age | 2.05E+00 | 1.22E-01 | 16.8 | 276 | 4.03E-44 | 1.81E+00 | 2.29E+00 |
|  | RH | Seg | 8.19E-02 | 9.93E-02 | 0.82 | 276 | 4.10E-01 | -1.14E-01 | 2.77E-01 |
|  | RH | Age x Seg | 4.60E-03 | 1.33E-02 | 0.34 | 276 | 7.31E-01 | -2.17E-02 | 3.08E-02 |
| IS | RH | Age | 1.71E+00 | 1.27E-01 | 13.39 | 276 | 7.34E-32 | 1.46E+00 | 1.96E+00 |
|  | RH | Seg | -4.24E-01 | 1.15E-01 | -3.7 | 276 | 2.57E-04 | -6.49E-01 | -1.99E-01 |
|  | RH | Age x Seg | 7.59E-02 | 1.54E-02 | 4.94 | 276 | 1.38E-06 | 4.56E-02 | 1.06E-01 |
| ML | RH | Age | 2.58E+00 | 1.26E-01 | 20.52 | 276 | 1.82E-57 | 2.34E+00 | 2.83E+00 |
|  | RH | Seg | 5.80E-01 | 1.20E-01 | 4.84 | 276 | 2.16E-06 | 3.44E-01 | 8.16E-01 |
|  | RH | Age x Seg | -1.39E-01 | 1.61E-02 | -8.67 | 276 | 3.80E-16 | -1.71E-01 | -1.08E-01 |

**Supplementary Figure 1.**
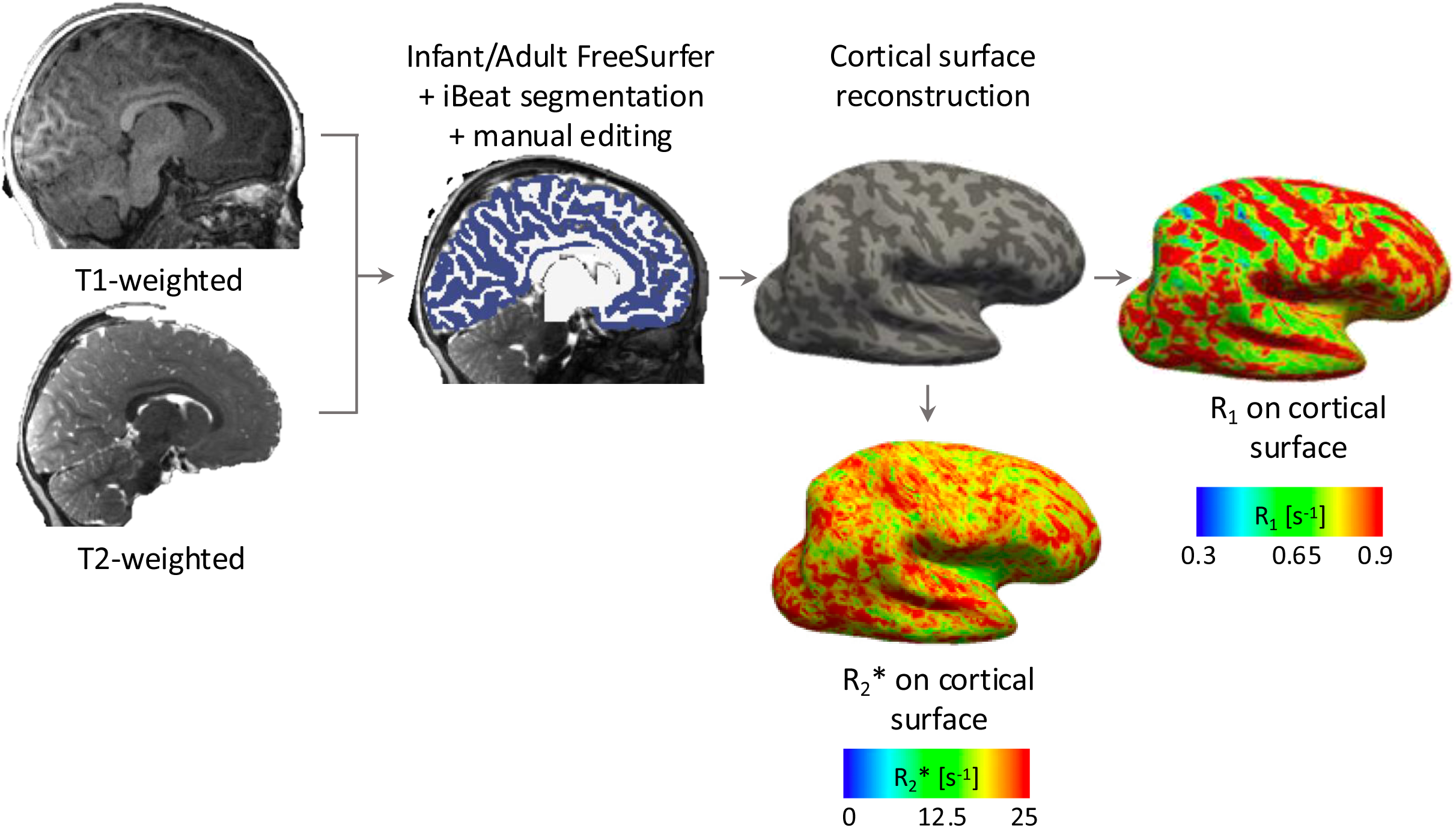
MRI data preprocessing pipeline and microstructural map. Schematic showing the preprocessing pipeline associated with obtaining the white and gray matter segmentations for generating the cortical surfaces (sample right hemisphere surface) obtained using infant or adult FreeSurfer’s automated pipeline and microstructural quantitative R_1_ [s^−1^] and R_2_* [s^−1^] maps aligned to the same anatomical brain volume and cortical surface. All analyses are done for each individual participant in their native brain space.

**Supplementary Figure 2.**
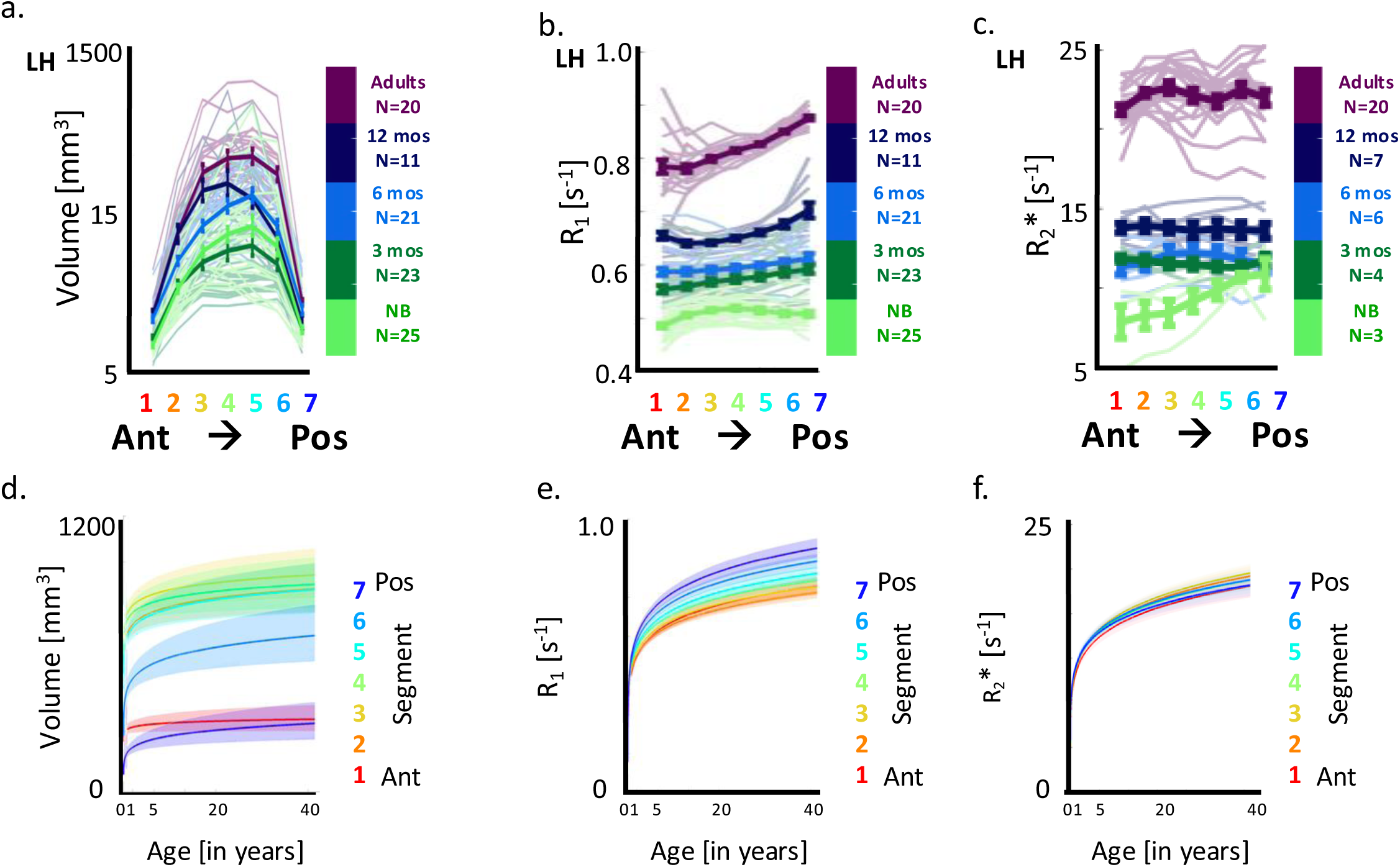
Spatial gradients in left putamen’s macro and microstructure along the anterior-posterior axis. a) Volume [mm^−3^] of the left putamen in infants and adults. Thinner lines: represent individual participant’s trend. Error bars: standard error across participants per age group. b-c) same as in a for R_1_ and R_2_*. d-f) Curves representing changes in volume, R_1_ and R_2_* as a function of age per segment. LH: left hemisphere.

**Supplementary Figure 3.**
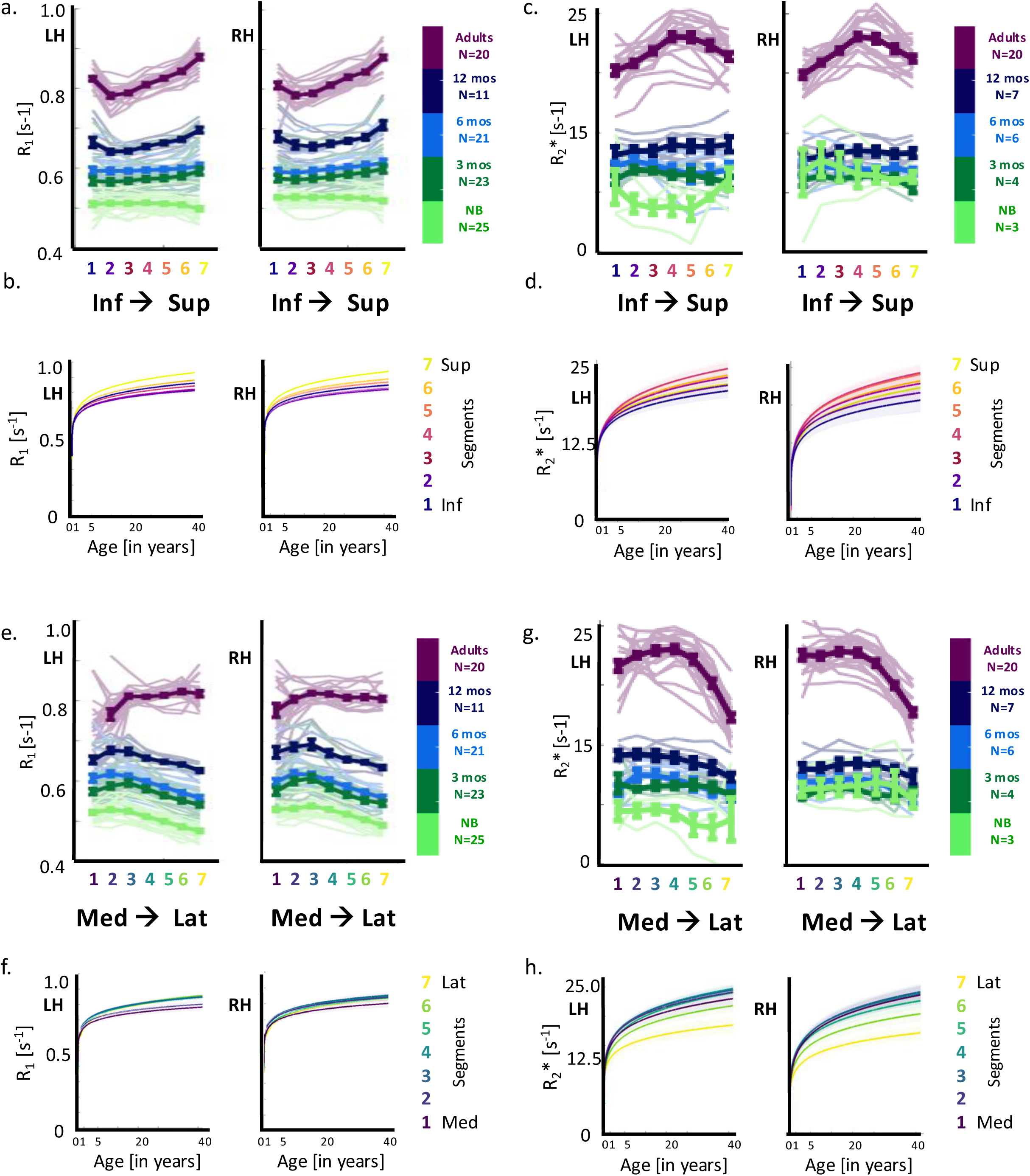
Development of microstrucural gradients in putamen from infancy to adulthood along its inferior-superior and medial-lateral axes in both hemispheres. a) R_1_ gradients along the inferior-superior (IS) axis in the left and right putamen of newborns, 3-month-, 6-month-, 12-month-old infants and adults. Error bars represent standard error across participants per age group. b) Curves representing R_1_ changes as a function of age per putamen segment. c-d) same as in a,b for R_2_*. e-h) Same as in a-d for medial-lateral axis. LH/RH: left/right hemisphere. Full statistics in Supplementary Tables 2-3.

**Supplementary Figure 4.**
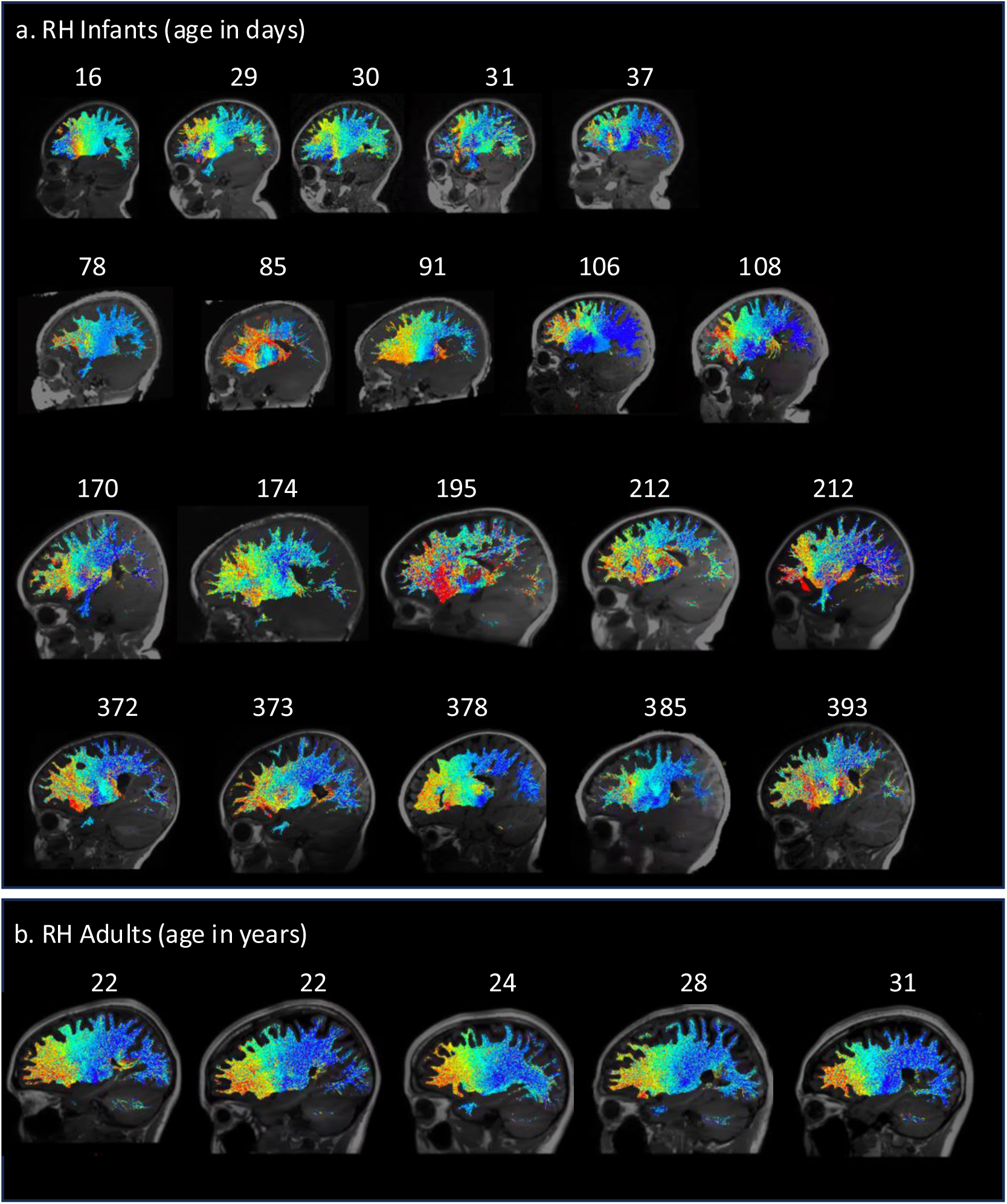
Cortico-putamen connections of the right putamen in sample infants across ages and adults. Sagittal images showing overlapping connectivity profiles of all seven segments of the right putamen along its anterior-posterior axis (colored from red to blue (A-P) in a) 5 sample newborns, 3-month-olds, 6-month-olds, 12-month-olds and b) 5 sample adults. Age in days for infants and years for adults are noted above each brain. Connections of each segment with cortex are colored respective to the color of the segment from anterior to posterior.

**Supplementary Figure 5.**
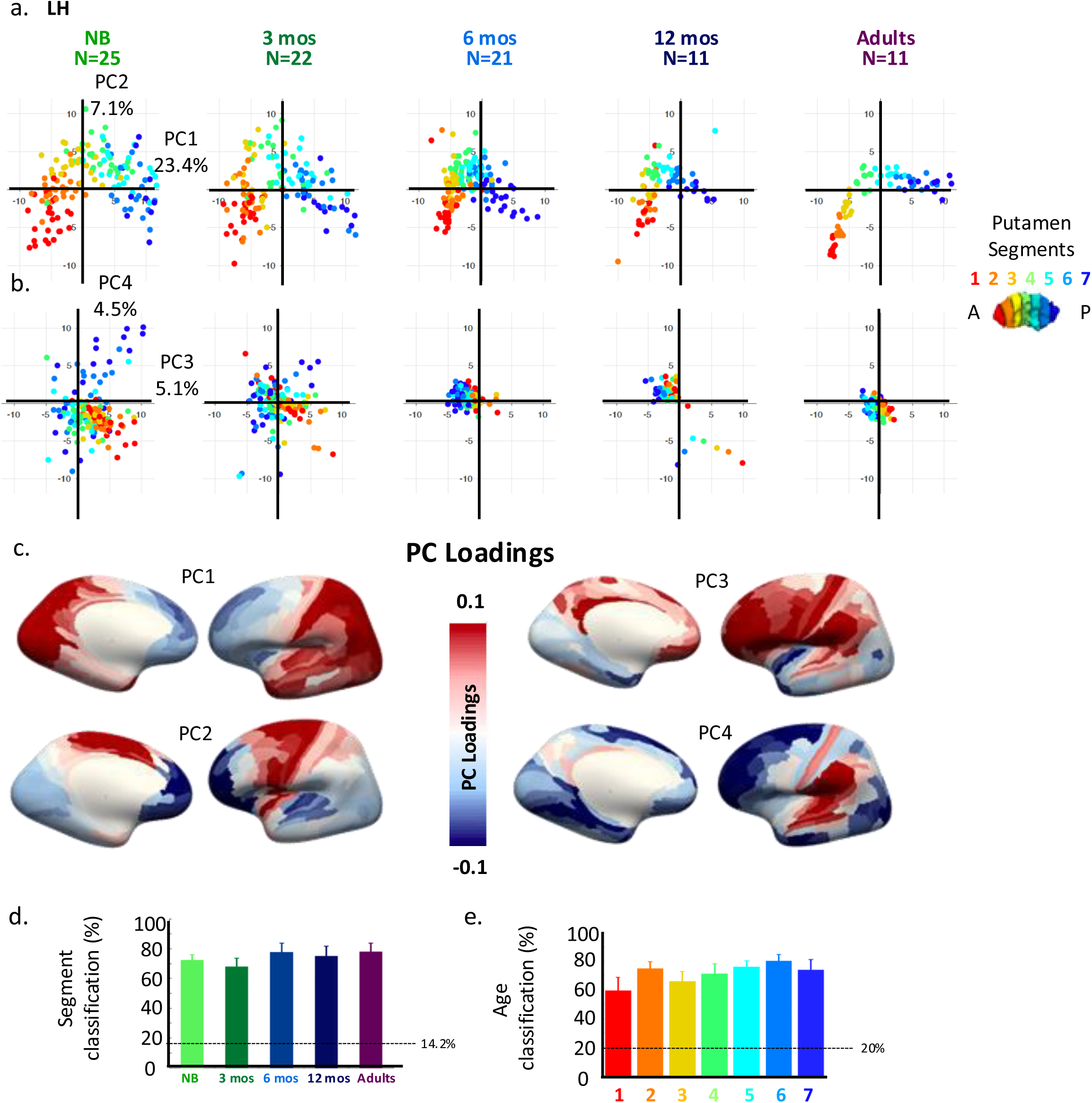
Principal component analysis on connectivity profiles of the left hemisphere showing CPs are organized by anterior-posterior segments of the left putamen and participant’s age. a) White matter connectivity profiles (CP) projected on the first two principal components (PC1 and PC2) colored by segment color (anterior to posterior: red to blue) per age group (N=5, (light green: newborns, dark green: 3-months-old (mos), light blue: 6 mos, dark blue: 12 mos, purple: adults). Each dot is a CP per segment. b) Same as in a for PC3 and PC4. c) Individual PC (PC 1-4) loadings projected onto average adult FreeSurfer brain surface delineated using 180 regions of the Glasser Atlas(*65*) (blues: negative loadings, reds: positive loadings). d-e) Classification accuracies for classifying segments and age groups. We utilized the information across PCs explaining more than 80% of variance, N_PCs_ = 34, Overall, both segments (%mean accuracy±standard deviation: RH: 73.63±2.03%, chance: 14.28%) and age (%accuracy RH: 71.69±3.99%, chance: 20%) can be classified significantly above chance LH: left hemisphere.

**Supplementary Figure 6.**
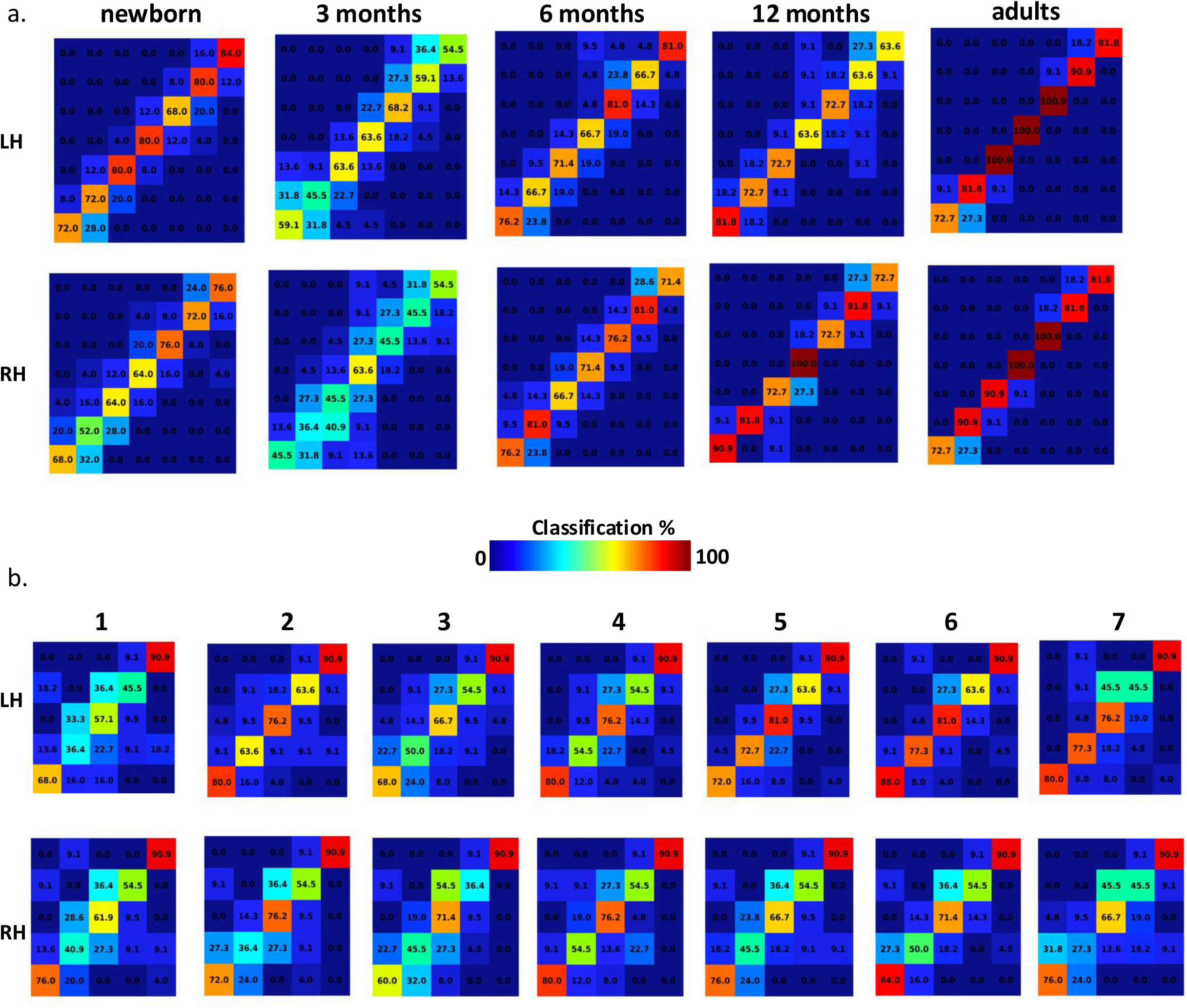
Confusion matrices showing classification accuracies for classifying segment and age from left and right putamen’s connectivity profiles. a) Classification accuracies for classifying left and right hemisphere segments using a multiclass model for support vector machines with leave-one-(participant)-out cross-validation, rows sum to 100 (Yellow: higher classification). b) same as in a for classifying age groups per segment (1-7: anterior to posterior). LH/RH: left/right hemisphere.

**Supplementary Figure 7.**
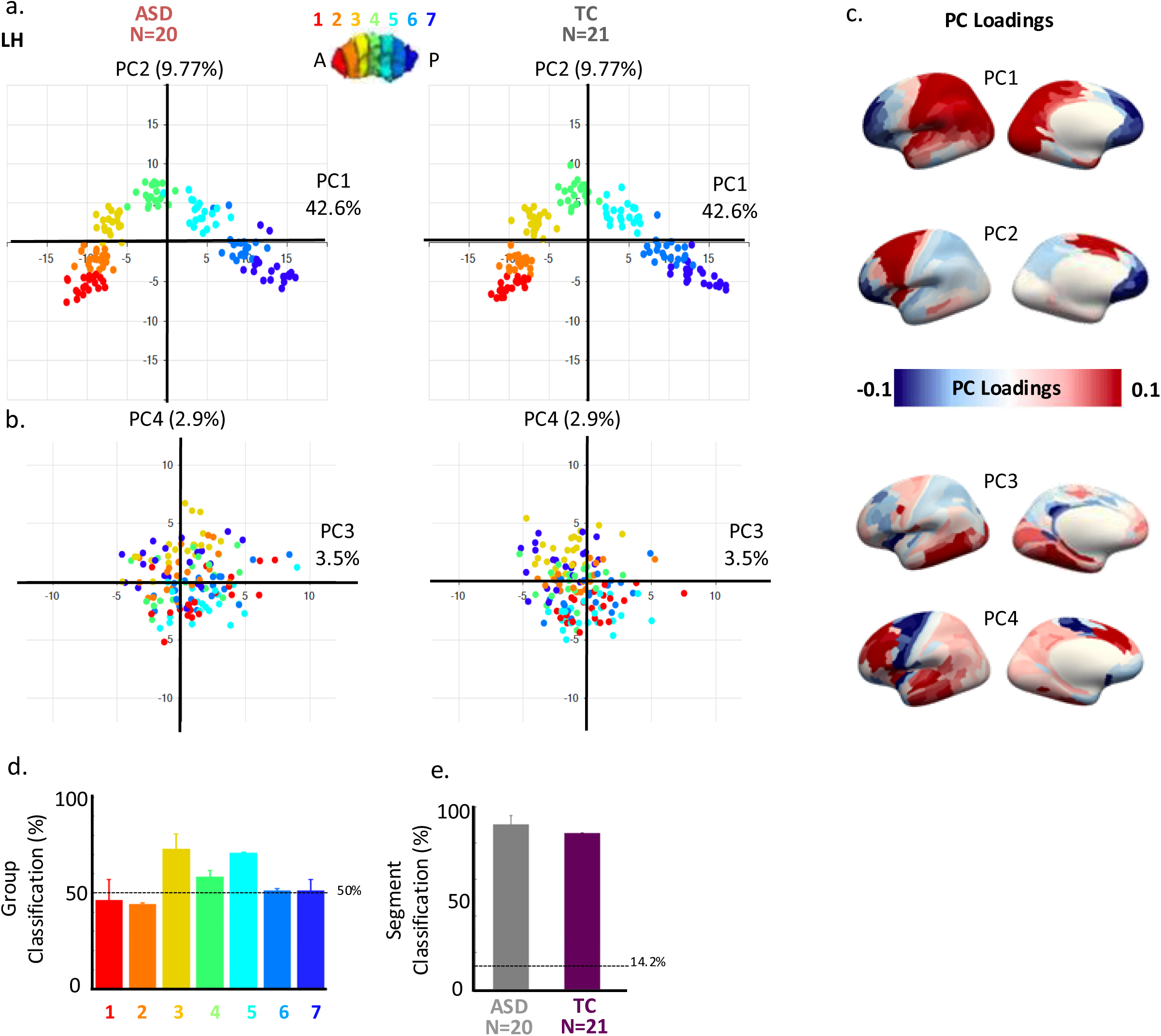
Cortico-putamen connectivity in ASD is organized like controls, with subtle overconnectivity with left superior temporal, and face processing areas in autism. a) Connectivity profiles (CF) of segments (N=7, left panel) and participants (N_ASD_=20, N_TC_=21) projected on PC1 and PC2. Each dot: CP per segment (red: anterior most to blue: posterior most) and participant (red: ASD; gray: typical controls). b) Same as in a) for PC3 and PC4. c) PC 1-4 loadings projected onto average adult FreeSurfer brain (blues: negative loadings, reds: positive loadings). d-e) Classification accuracy for classifying segments and groups from CPs. We utilized information across PCs explaining more than 58% of variance, N_PCs_ = 4. Segments (%accuracy LH: 94.04±0.61%, chance: 14.29%) were classified above chance however, group was classified with highest accuracy in segment 3 (%accuracy LH: 72.97±3.68 %, chance: 50%). LH: left hemisphere.

## Notes

### Competing Interest Statement

The authors have declared no competing interest.

